# Oct1 cooperates with Smad transcription factors to promote mesodermal lineage specification

**DOI:** 10.1101/2020.12.01.406488

**Authors:** Jelena Perovanovic, Yifan Wu, Hosiana Abewe, Zuolian Shen, Erik P. Hughes, Jason Gertz, Mahesh B. Chandrasekharan, Dean Tantin

## Abstract

The transition between pluripotent and tissue-specific states is a key aspect of development. Understanding the pathways driving these transitions will facilitate the engineering of properly differentiated cells for experimental and therapeutic uses. Here, we showed that during mesoderm differentiation, the transcription factor Oct1 activated developmental lineage-appropriate genes that were silent in pluripotent cells. Using mouse embryonic stem cells (ESCs) with an inducible knockout of Oct1, we showed that Oct1 deficiency resulted in poor induction of mesoderm-specific genes, leading to impaired mesodermal and terminal muscle differentiation. Oct1-deficient cells exhibited poor temporal coordination of the induction of lineage-specific genes and showed inappropriate developmental lineage branching, resulting in poorly differentiated cell states retaining epithelial characteristics. In ESCs, Oct1 localized with the pluripotency factor Oct4 at mesoderm-associated genes and remained bound to those loci during differentiation after the dissociation of Oct4. Binding events for Oct1 overlapped with those for the histone lysine demethylase Utx, and an interaction between Oct1 and Utx suggested that these two proteins cooperate to activate gene expression. The specificity of the ubiquitous Oct1 for the induction of mesodermal genes could be partially explained by the frequent coexistence of Smad and Oct binding sites at mesoderm-specific genes and the cooperative stimulation of mesodermal gene transcription by Oct1 and Smad3. Together, these results identify Oct1 as a key mediator of mesoderm lineage–specific gene induction.

## INTRODUCTION

Lineage specification is the key process in the development of multicellular organisms by which undifferentiated cells progressively acquire tissue-and cell type-specific features (*1*). It is dynamically regulated, requiring extensive transcriptional, epigenetic, and topological remodeling to selectively activate lineage-appropriate gene expression programs and stably repress the expression of genes specific for alternative lineages. Embryonic stem cells (ESCs) represent a pluripotent cell type capable of both self-renewal and differentiation into all three embryonic germ layers (*2*). The three germ layers are established during gastrulation – a spatial reorganization of the embryo from a single-layer epiblast into the multilayered post-implantation embryo. One of the germ layers – mesoderm (MD) – gives rise to dermomyotome (muscle and dermis), sclerotome (axial skeleton), and lateral MD (cardiac) among other tissue types.

The transcriptional changes underlying lineage specification require extensive chromatin remodeling and spatiotemporal activation of genes encoding developmental master regulators. Remodeling is potentiated by a unique chromatin landscape in pluripotent cells. Chromatin in ESCs is largely accessible and lacks stable heterochromatin domains (*3, 4*). A large number of genes encoding lineage-specific developmental regulators are marked at promoters and gene bodies by covalently modified nucleosomes that simultaneously convey activating (H3K4me3) and repressing (H3K27me3) potential (*5, 6*). In ESCs, these “bivalent” genes are silent or expressed in low amounts but lack DNA methylation and are poised for activation. During development, gene bivalency is thought to resolve either by removal of activating marks and gene silencing, or removal of repressive marks and gene activation. Which poised genes resolve in one direction or the other is lineage-specific, resulting in distinct, cell fate–appropriate transcriptional programs and durable repression of lineage-inappropriate genes.

POU domain–containing transcription factors play central roles in development (*7*). The POU factor Oct4 (encoded by *Pou5f1*) is a master regulator of the induction and maintenance of pluripotency (*8–10*). Oct4 associates with bivalent genes in ESCs (*5*), and can briefly pair with lineage-specifying transcription factors early in development as pluripotency is lost and cells form mesendoderm (*11–14*), but is silenced in their differentiated progeny before bivalency resolution and the induction of tissue-and cell type-specific gene expression (*14*). A second POU protein, Oct1 (also known as Pou2f1) is co-expressed with Oct4, but unlike Oct4 is expressed beyond pluripotency (*15*). Oct1 is widely expressed and in mice and is required for placental and embryonic development (*16, 17*). Circumventing the placental defects due to the loss of Oct1 by using tetraploid complementation results in developmental arrest at embryonic day 8.25 (E8.25) with no more than five somites (*15*). Oct1-deficient ESCs are morphologically normal and proliferate normally, but upon differentiation show decreased developmental lineage-specific gene expression and increased expression of developmentally inappropriate genes, including genes normally expressed only in extra-embryonic tissues (*15*). Extended incubation of these cells supports an interpretation that there is a failure to differentiate rather than a kinetic delay (*15*).

Here, using Oct1 inducible-conditional knockout mouse ESCs, we show that pluripotent cells lacking Oct1 failed to form terminally differentiated myotubes, but Oct1 retroviral complementation restored differentiation. Bulk RNA sequencing (RNA-seq) profiling early in the differentiation timecourse identified gene expression abnormalities early in MD differentiation that predicted the later phenotypic defects. Single-cell RNA-seq (scRNA-seq) revealed that cells lacking Oct1 accumulated abnormal cell populations associated with epithelial characteristics, oxidative stress, and early MD identity, while almost completely failing to reach somite-stage differentiation. Pseudotime analysis revealed increased predilection to proceed down incorrect developmental trajectories, and loss of coherence in differentiation gene expression programs. Oct1 and Oct4 chromatin immunoprecipitation sequencing (ChIP-seq) identified co-binding of Oct1 with Oct4 in pluripotent cells that was carried forward into differentiation after Oct4 was lost. H3K27Ac chromosome conformation capture coupled to chromatin immunoprecipitation (Hi-ChIP) revealed that a large number of topological chromatin interactions that were gained with differentiation corresponded to sites of Oct1 binding. Oct1 interacted with components of the Utx (also called Kdm3a) H3K27me3 demethylase complex, and its binding strongly overlapped with that of Utx at developmentally appropriate target genes, consistent with removal of H3K27me3 and derepression of tested targets. Binding sites for Oct1 and Oct4 in pluripotent cells in which Oct1 binding was carried forward during MD differentiation were also enriched in binding sites for Smad transcription factors, key mediators of MD specification. Oct1 and Smad3 interacted and cooperatively activated gene expression, providing a means of restricting Oct1’s activating potential to MD-specific genes. The cumulative results support a model in which Oct1 “canalizes” differentiation by supporting the induction of lineage-specific genes.

## RESULTS

### Loss of Oct1 causes aberrant mesodermal differentiation

To study the consequences of Oct1 deficiency for MD development, we subjected Oct1 inducible-conditional *Pou2f1^fl/fl^*;Rosa26-CreER^T2^;LSL-YFP control mouse ESCs (hereafter, “parental” cells) and tamoxifen-treated, Oct1-deficient cells derived from them (*15*) (hereafter, “cKO”) to an MD differentiation protocol that begins to generate myotubes after >10 days of culture. The differentiation process is initiated by bone morphogenetic protein 4 (Bmp4) treatment to activate transforming growth factor-β (TGF-β) signaling and generates dynamic changes in metabolic programs (*18*), epithelial-to-mesenchymal transition (EMT) (*19*), and induction of MD-specific gene expression programs during the differentiation timecourse (*20*). Parental and cKO cells were differentiated for 11 days (D11) to generate early myotubes.

Immunostaining revealed that embryonic myosin heavy chain (MyH-emb) expression was robust in fused myotubes from parental cells but close to background in cKO cells (Fig.1A). We then queried expression of myogenic genes (*Myod* and *Myog*) using more fully differentiated parental and cKO cells (D19) and RT-qPCR. Relative to *Rps2* (which encodes a ribosomal 40S subunit), both genes were strongly expressed in parental but not cKO cells (Fig.1B). These results demonstrate that cKO ESCs induced to differentiate into MD manifest defective early myogenic gene expression and fail to differentiate into muscle lineages.

**Fig. 1.**
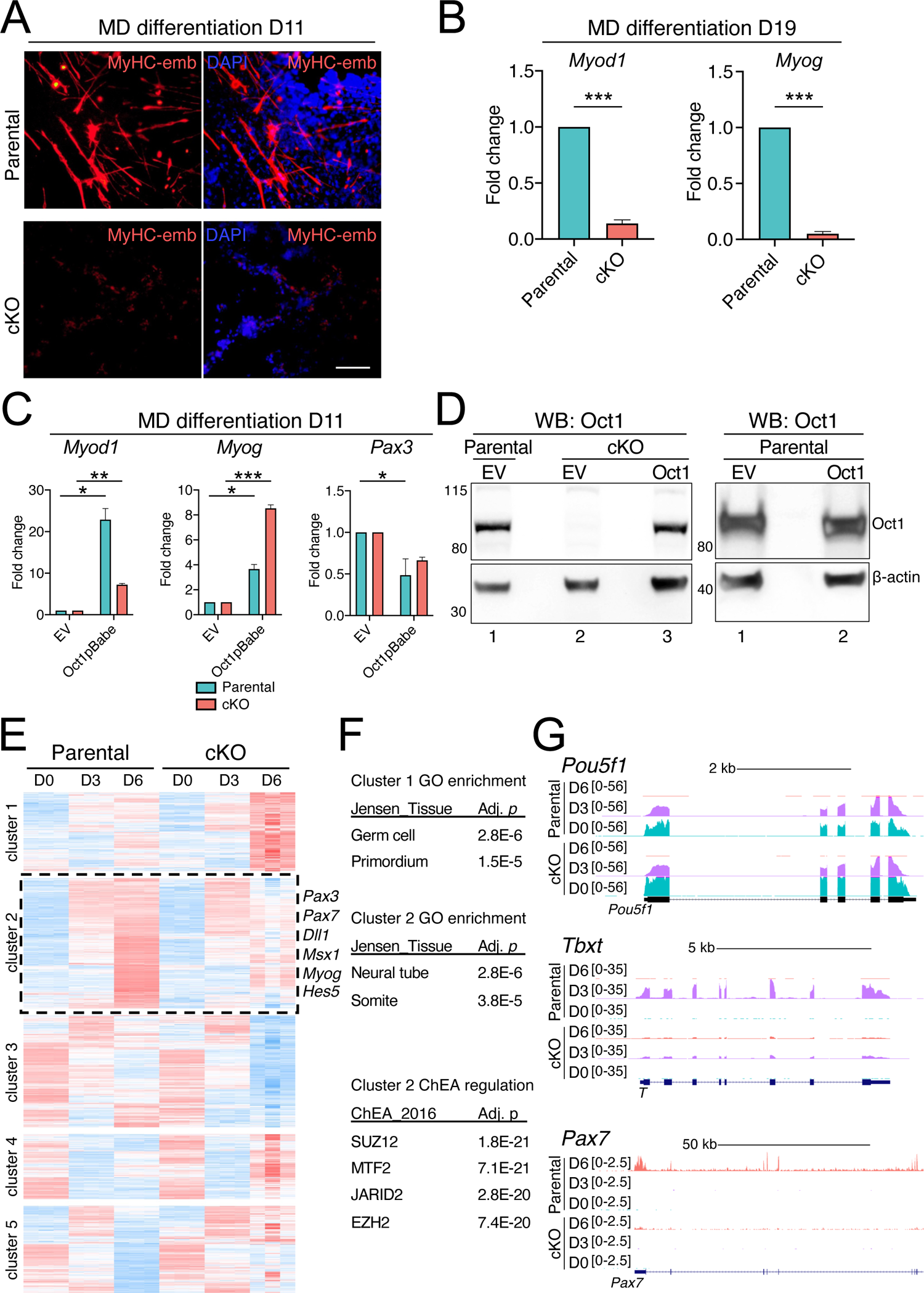
Failed MD and muscle lineage induction in differentiating Oct1 cKO ESCs. (**A**) Embryonic myosin heavy chain (MyHC-emb) expression alone and merged with DAPI is shown in parental and Oct1 cKO cells at MD differentiation day 11 (D11). Nuclei are labelled with DAPI. Scale bar, 100 μm. Images are representative of 3 independent experiments. (**B**) Relative expression of the myogenic genes *Myod1* and *Myog* in parental and Oct1 cKO ESCs differentiated for 19D. Data represent an average of N=3 biological replicate experiments. Error bars depict ±SEM. (**C**) RT-qPCR for the myogenic genes *Myod1*, *Myog*, and *Pax3* in parental and cKO cells transduced with retroviruses encoding Oct1 or empty vector (EV). Transduced cells were selected using puromycin for 48 hr prior to differentiation for 11D. EV values were set to 1. Data represent an average of N=3 biological replicates. Error bars depict ±SEM. (**D**) Immunoblot showing ectopic Oct1 expression in parental and cKO cells. β-actin is a loading control. Numbers to the left of the blots note positions of molecular weight markers in kilodaltons. Blots are representative of 2 independent experiments. (**E**) Bulk RNA-seq heatmap of differentially expressed genes (parental vs. cKO) at D0, D3, and D6 of MD differentiation is shown. N=3 biological replicates were performed per condition. Representative genes showing poor induction in cKO cell cluster 2 are noted. (**F**) “Jensen_Tissue” GO enrichment for the differentially regulated genes in Clusters 1 and 2 in (B), and ChIP-X Enrichment Analysis (ChEA) query results for the differentially regulated genes in Cluster 2 in (B), are shown. Adj. *p*, adjusted *p*-value. (**G**) Example RNA-seq genome tracks (*Pou5f1*, *Tbxt*, *Pax7*) are shown. *Pax7* is an example cluster 2 gene. The indicated gene, with intro/exon structure, is show at the bottom of each set of tracks. Track height indicates the density of bulk RNA-seq reads at that genomic position. Teal indicates D0, purple indicates D3 and red indicates D6. Scale is shown at top for each set of tracks. kb, kilobases. Y-axes were group-autoscaled for each gene.

To determine the effect of ectopic Oct1 expression, we transduced parental and cKO ESCs with retroviral vectors encoding murine Oct1 or empty vector (EV) controls. The vectors encode a puromycin resistance cassette, allowing for selection of transduced cells. Immediately after selection, populations of transduced cells were subjected to MD differentiation to D11, and tested for *Myod*, *Myog*, and *Pax3* mRNA expression by RT-qPCR. Differentiating cKO ESCs transduced with Oct1 expressed *Myod* and *Myog* more strongly. In contrast, *Pax3* expression was significantly reduced in cKO cells (Fig.1C). The combination of increased *Myod* and *Myog* with diminished *Pax3*, a transiently expressed inducer of the myogenic program, at late differentiation timepoints suggests that restoration of Oct1 allows cells to more efficiently transit through a *Pax3*-expressing intermediate state, such that more cells enter into a terminal myogenic program. Retrovirally expressed ectopic Oct1 expression also increased expression of terminal muscle differentiation genes in parental cells, with a trending reduction in *Pax3* expression (Fig.1C). Immunoblotting confirmed ectopic Oct1 expression and complementation of Oct1 deficiency (Fig.1D).

### Lineage-specific gene expression is defective in differentiating Oct1-deficient cells

To identify how Oct1 loss perturbs MD differentiation in greater detail, we performed bulk RNA-seq using parental and Oct1 cKO cells. We identified ∼1700, ∼800, and 3,300 significantly differentially expressed genes at D0, D3 and D6, respectively (FDR ≤ 0.05; −1 < log2FC < 1, table S1). Euclidean distance analysis revealed tight correlations between replicates and between parental and cKO cells at D0 and D3, but divergence at D6 relative to the other conditions and between parental and cKO cells at D6 (fig.S1). The D0 gene expression changes, which were selected to be over 2-fold, were smaller in magnitude, with D0 genes changed by an average of 2.7-fold (down) or 3.3-fold (up), vs. D6 genes 3.4 fold (down) and 3.9-fold (up, table S1). D0 changes were also associated with unremarkable gene ontology (GO) terms such as “ion transmembrane transporter activity” that were associated with weak *p*-values. One notable gene with increased abundance in D0 Oct1 deficient ESCs was *Pou2f3*, which encodes Oct11.

Unsupervised hierarchical clustering revealed groups of genes regulated similarly between parental and cKO cells, and others with differential expression (Fig.1E). Cluster 2, for example, showed strong induction in differentiating parental cells but failed induction in cKO cells at D6. Example genes in this set include *Pax3*, *Pax7*, and *Myog* (Fig.1E). GO analysis of the differentially expressed genes in cluster 2 using “Jensen_Tissue” categories revealed association with differentiation to neural tube (neuroectoderm) and somite (Fig.1F). By contrast, clusters 1 and 4, which showed aberrant gene expression in differentiating cKO cells, were associated with lineage-inappropriate Jensen_Tissue terms such as primordial germ cells (Fig.1F). To identify potential regulators of cluster 2 genes with failed induction in differentiating cKO cells, we queried the ChIP Enrichment Analysis (ChEA) database (*21*). The genes in cluster 2 tended to also be bound by the polycomb repressor complex 2 (PRC2) components Suz12 and Mtf2 (Fig.1F). Example tracks are shown in Fig.1G for *Pou5f1* (pluripotency), *Tbxt* (also called *T* or *Brachyury*, cluster 3), and *Pax7* (cluster 2). The retained D3 expression of *Pou5f1*, which encodes Oct4, in both parental and cKO cells provides a likely explanation for the tight correlations between D0 and D3 cKO vs. parental gene expression states, with divergence at D6 (fig.S1), because the presence of Oct4 may mask the loss of Oct1 at D3. These data indicate that Oct1 loss results in defective induction of developmental genes that are also regulated by chromatin-modifying activities that act on H3K27me3.

### Differentiating Oct1-deficient ESCs proceed down abnormal developmental trajectories

To investigate perturbations in cell populations and gene expression at single-cell resolution, we performed single-cell RNA-seq (scRNA-seq) using parental and cKO ESCs, and cells differentiated towards MD for 3 or 6 days. Cell numbers ranged between 1000 and 2500, with reads per cell ranging between 90,000 and 200,000. Data from the different conditions were subjected to graph-based clustering and uniform manifold approximation and projection (UMAP) to visualize clusters of cells with similar gene expression. Integrated analysis of parental cells in a pluripotent state (D0) at the two differentiation timepoints identified a range of expected cell populations, from naïve and primed pluripotency to neuromesodermal, paraxial MD, and early somite (Fig.2A). These populations recapitulate developmental stages associated with early MD (*Fgf17* and *Tbxt*/*T*/*Brachyury*), neuromesoderm (*Hoxc9*, *Pax3*, *Sox2*), paraxial MD (*Meox1*, *Twist1*, *Pax3*), and early somite and neuronal progenitors (*Cxcr4, Uncx*) (Fig.2B, fig.S2). These results provide a single cell–resolution map of early MD differentiation.

**Fig. 2.**
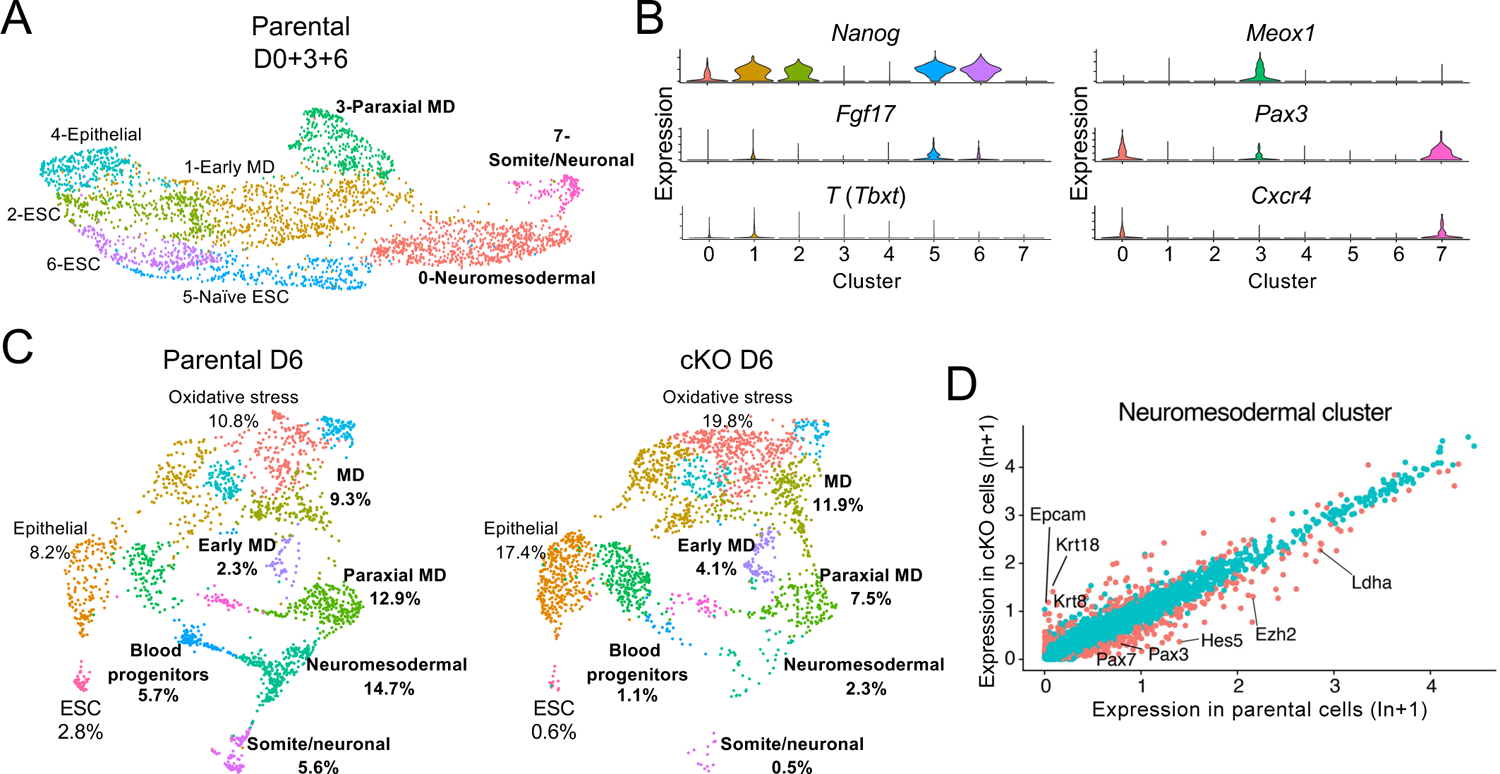
Differentiating Oct1-deficient ESCs canalize poorly into mesodermal lineages. (**A**) UMAP projection of scRNA-seq data from superimposed parental (Oct1-sufficient) undifferentiated ESCs (day 0, D0), and parental cells early during MD differentiation (D3 and D6). Clusters of cells were labeled based the expression of developmental markers as in (B). MD-associated clusters are shown in bold. N=3 combined biological replicate plates were used for the scRNA-seq. (**B**) Violin plots showing expression of key developmental markers by cluster. Data were log-normalized for each cell using the natural logarithm, scaled, and averaged using mean(expm1(x)). (**C**) Comparative UMAP projections of integrated D6 parental and Oct1-deficient (cKO) scRNA-seq populations. Clusters were labeled computationally and identified based on gene expression. Relative frequencies are shown. MD lineage–associated clusters are shown in bold. (**D**) Differential gene expression analysis of the neuromesodermal cluster shown as a scatter plot. Red dots depict significantly differentially expressed genes based on FDR corrected *p*< 0.05 and fold change >1.2. Example differentially expressed genes are labeled.

We compared the parental controls to cKO cells to identify changes in cell populations and gene expression. In agreement with the bulk RNA-seq data, D0 and D3 cKO cells showed few differences from parental, Oct1-sufficient cells. In contrast, D6 cKO cells showed poor differentiation capacity, with reductions in key mesodermal clusters (Fig.2C, table S2). In parental cells, populations characterized by neuromesodermal and paraxial MD gene expression accounted for 14.7 and 12.9% of cells, respectively, whereas populations associated with blood progenitors and somites accounted for 5.7% and 5.6% of cells, respectively (Fig.2C). The complexity in the populations is consistent with findings that somites are derived from multiple transcriptional trajectories including early paraxial mesoderm and neuromesodermal progenitors (*22*). In contrast, cKO D6 cells showed increases in clusters that retain epithelial characteristics and strongly decreased percentages of neuromesodermal progenitors (2.3%, a six-fold decrease), paraxial MD (7.5%, two-fold), blood progenitors (1.1%, >5-fold), and somites (0.5%, >10-fold, Fig.2C, right panel). Gene expression comparisons between parental and cKO paraxial mesoderm clusters revealed that cKO cells failed to robustly induce lineage-appropriate genes such as *Pax3* and *Pax7*, and inappropriately increased the expression of lineage-inappropriate markers such as the epithelial-specific genes *Krt8* and *Krt18* (Fig.2D). These results show that Oct1 is necessary for accurate MD differentiation and to suppress expression of genes for alternative lineages.

Pseudotime analysis of scRNA-seq data allows multiple differentiation timepoints to be overlaid with defined starting (ESC) and endpoint (somite) populations. Applying pseudotime, we found that parental control cells progressed through a largely linear pathway with one minor branch retaining an inappropriate epithelial fate (Fig.3A). In contrast, differentiating cKO ESCs showed a large proportion of cells inappropriately branching into alternative developmental trajectories (Fig. 3A), consistent with the diminished developmental progression of paraxial MD to the somite stage, and consistent with enrichment of cells that inappropriately maintain an epithelial state (Fig.2C, fig.S2). We also examined pseudotemporal gene expression using specific genes associated with pluripotency and MD development. In this analysis, the position of each cell in pseudotime is shown on the X-axis and the degree of expression for the indicated gene on the Y-axis (Fig.3B). Parental cells showed robust early expression of genes associated with pluripotency, such *as Klf4*, which was lost roughly mid-way through pseudotime. *Tbxt* was transiently induced, whereas the early somite markers *Pax7* and *Cxcr4* were induced later (Fig.3B). In contrast, cKO cells exhibited prolonged expression of *Tbxt* and largely failed to increase expression of *Pax7* and *Cxcr4* (Fig. 3B). We used pseudotemporal ordering to visualize expression of the 2000 most variable (dynamic) genes in parental cells and compared expression of the same genes in the same order in cKO cells (Fig.3C). This analysis revealed clusters associated with pluripotency (*Klf4*, *Nanog, Pou5f1*), early MD differentiation (*Tbxt*), paraxial mesoderm (*Rspo3, Meox1*), and early somites (*Dll1, Pax3*). The pseudotime gene expression pattern for cKO cells revealed largely normal expression of early gene expression markers but loss of temporal coherence at subsequent steps. These “fuzzy” gene expression patterns began early in differentiation, with, for example, prolonged expression of the MD master regulator *Tbxt* and noncoherent expression of genes associated with paraxial mesoderm such as *Meox1* and *Rspo3*. The diffuse pattern was different from what would be expected with a kinetic delay, which would cause a rightward shift. In addition, the induction of somitic genes normally expressed later in development, such as *Dll1*, *Pax3*, and *Pax7*, was poor in cKO cells (Fig.3C).

**Fig. 3.**
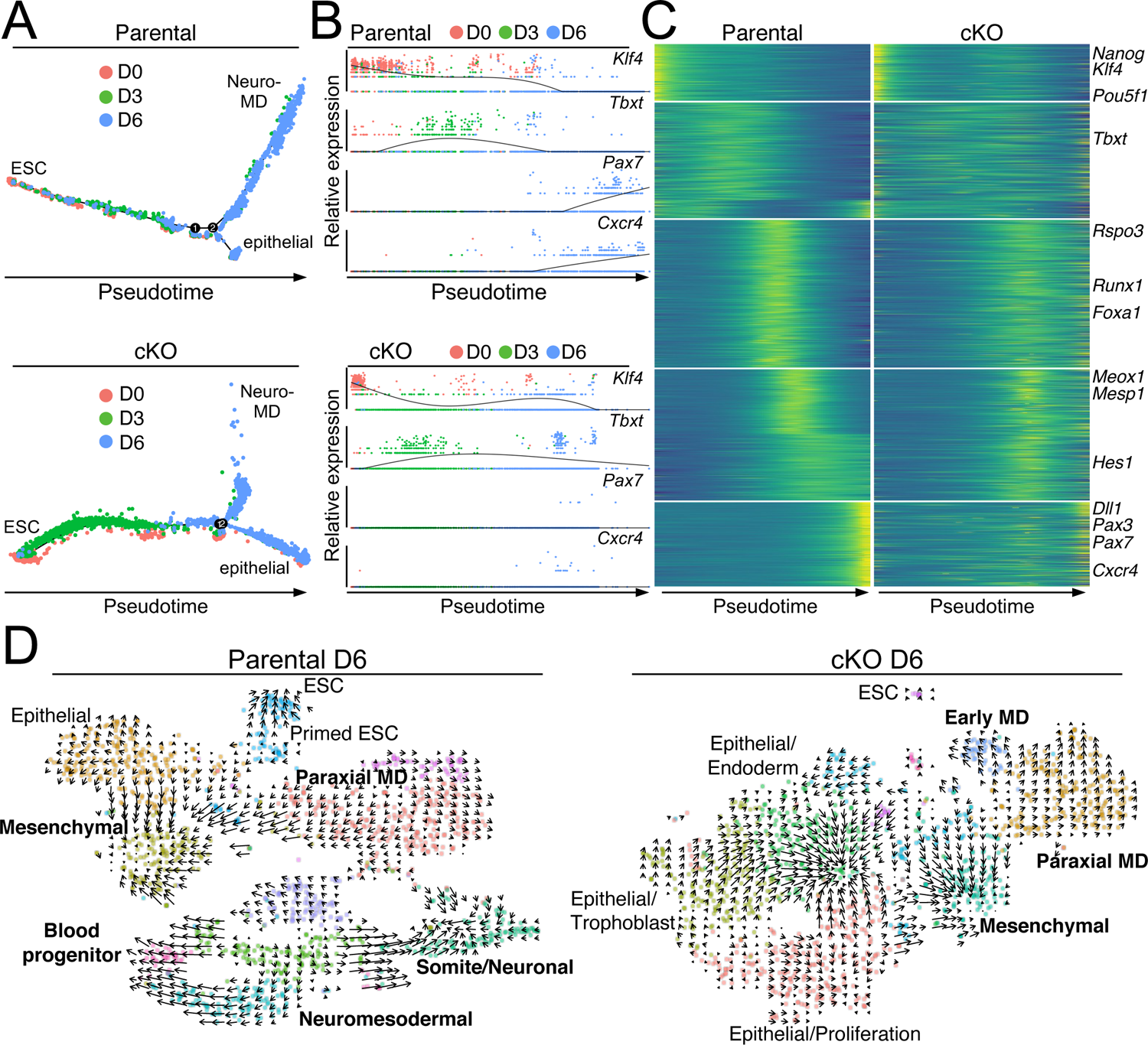
Oct1-deficient cells show perturbed developmental trajectories. **(A)** Pseudotime analysis of pluripotent (D0) and differentiating (D3, D6) parental and cKO cells. Colors correspond to the time point at which cells were collected. (**B)** *Klf4*, *Tbxt*, *Pax7*, and *Cxcr4* mRNA expression across pseudotime in parental and cKO cells at D0, D3, and D6. Black trendline represents an average expression for a given gene across all populations. (**C)** Heatmap depicting expression of the 2000 most dynamically expressed genes (based on FindVariableFeatures function, Seurat) in parental D6 cells. Gene expression was plotted as a heat map across pseudotime in parental and cKO cells. Dynamic genes were first hierarchically clustered in parental cells to cluster groups of genes that behave similarly in pseudotime, then plotted in the same order in cKO cells. (**D)** Velocity gene expression analysis of parental and cKO cells at D6. Arrows point toward cells with gene expression closest to the future state of each cell based on unspliced/spliced transcripts. Arrow length represents magnitude. Clusters labelled in bold are associated with MD differentiation.

RNA velocity analysis allows developmental directionality to be inferred relative to other cells by comparing the ratio of unspliced, newly-synthesized pre-mRNA to mature, spliced mRNA. A vector is then assigned to each cell that indicates developmental direction and magnitude relative to the other cells. We clustered the cells separately by genotype (Fig. 3D), resulting in different patterns compared to cells clustered together in the UMAP analysis (Fig. 2C). Applying velocity to this clustering, we found that D6 differentiated parental cells showed largely ordered and canonical developmental progression, for example, paraxial MD-to-somite (Fig.3D). cKO cells by contrast were marked by convergent or stationary epithelial profiles indicative of failed differentiation potential, with cells progressing primarily towards a poorly differentiated state characterized by multiple lineage markers such as intestine and trophoblast (Fig.3D, left side epithelial clusters). Cumulatively, the data demonstrate that Oct1 is required for efficient early adoption of the MD differentiation program, with Oct1-deficient cells unable to canalize appropriate lineage specification programs.

### Oct1 occupies developmentally-regulated genes in ESCs

One model that explains the above findings is that Oct1 occupies developmental-specific targets bound by Oct4 in ESCs to mediate induction of developmentally appropriate genes and repression of genes specific to alternative lineages. To test this hypothesis, we performed ChIP-seq using antibodies against Oct1 and Oct4. Prior work showed that Oct1 occupancy at Oct4 targets increases during retinoic acid (RA)-mediated differentiation when Oct4 expression is lost (*15*). RA ultimately differentiates cells into neuroectodermal lineages (*23*). In undifferentiated ESCs, strong Oct1 binding (with Oct4) was only observed in a group of ∼100 genes containing Oct protein variant binding sites termed MOREs (more-palindromic octamer recognition element) (*15*). Here, we used a different Oct1 antibody with superior enrichment properties to perform ChIP-seq with undifferentiated parental Oct1-sufficient ESCs, as well as cells differentiated towards MD for 3 or 6 days. Oct4 ChIP-seq in undifferentiated ESCs (D0) was performed as a parallel control (fig. S3, A to C).

In pluripotent cells (D0), ∼22,000 Oct4 peaks were identified, corresponding to ∼6,000 genes with transcription start sites (TSS) within 20 kb (table S3). We found that ∼45% of Oct4 targets directly overlapped (0 bp) Oct1 peaks. Conversely, ∼60% of Oct1 targets directly overlapped Oct4 peaks (Fig.4A), indicating substantial overlap between binding of the two transcription factors. Shared Oct1 and Oct4 targets in ESCs included *Polr2a*, which encodes the largest subunit of RNA polymerase II, *Pou5f1*, which encodes Oct4, and *Dll1*, which encodes a developmentally-inducible mediator of Notch signaling expressed in the MD lineage, where it regulates muscle development (*24*). We also confirmed Oct1 and Oct4 binding to *Polr2a*, *Pou5f1* and *Dll1* using ChIP-qPCR (fig.S3, A to C). Oct1 binding to *Polr2a*, which contains two adjacent MOREs that can bind four Oct proteins (*25*), is exceptional relative to other genes in that it is far stronger than that of Oct4 (100× stronger for *Polr2a*, 3-10× weaker for *Pou5f1* and *Dll*). Re-ChIP (sequential ChIP) indicated that Oct1 and Oct4 bind these sites simultaneously. The signal was lost in Oct1 cKO ESCs, indicating specificity for Oct1 (fig.S3, A to C). Cumulatively, the data indicate that in ESCs Oct1 co-binds with Oct4 to an array of targets, including developmental-specific targets.

**Fig. 4.**
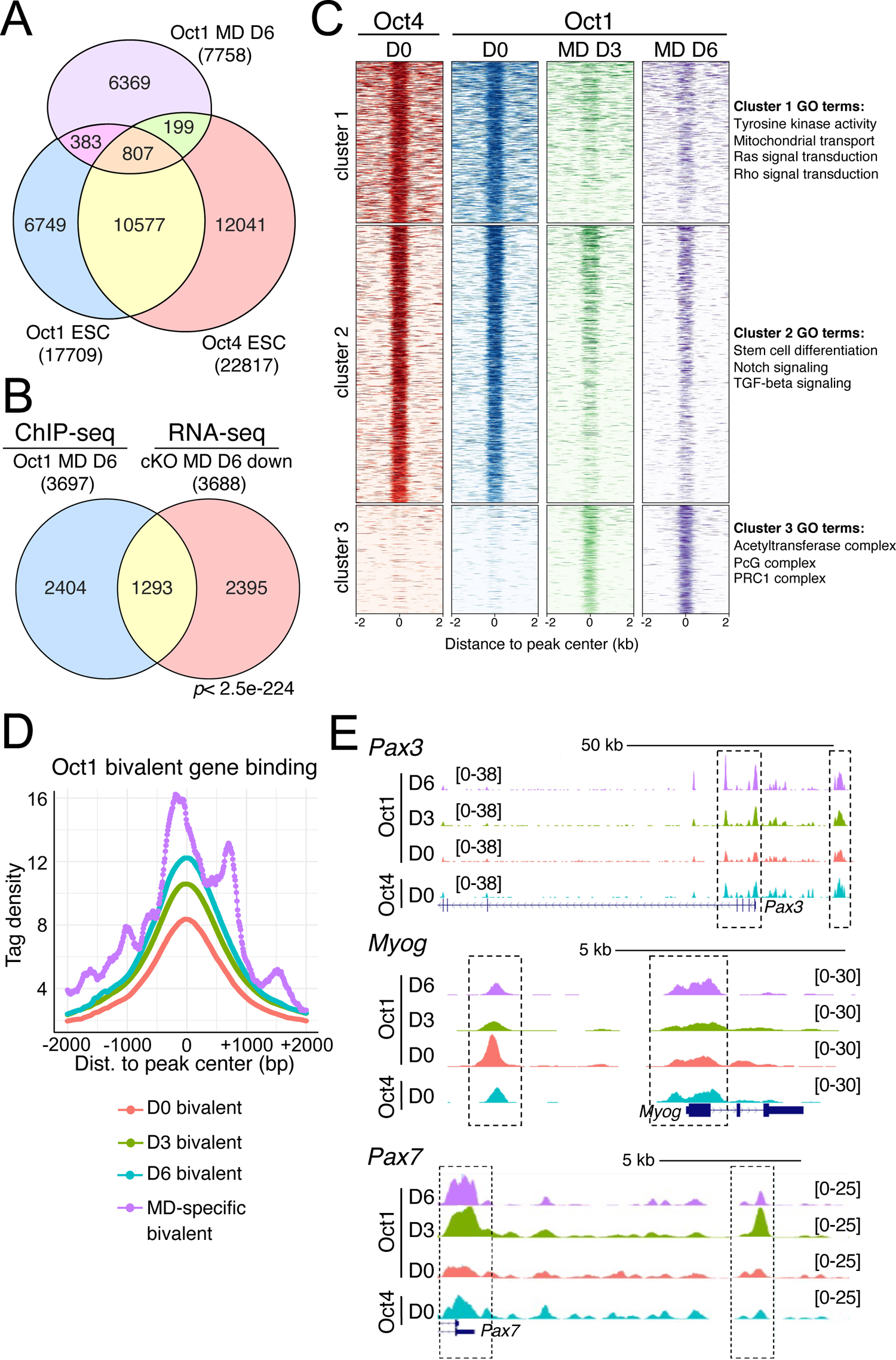
Oct1 co-occupies target sites with Oct4 in ESCs and continues to bind lineage-specific MD genes during differentiation. **(A)** Venn diagram showing common and unique Oct4 and Oct1 binding sites based on ChIP-seq in parental ESCs and at D6 of mesodermal differentiation. n = 3. (**B)** Comparison of Oct1 binding events (ChIP-seq) and gene expression (RNA-seq) reduced by Oct1 loss at MD differentiation day 6. *P*-value was computed using a hypergeometric test. Target genes (< 20 kb) were used rather than peaks, explaining the differing numbers compared to panel (A). (**C)** A matrix of Oct4 and Oct1 ChIP-seq enrichment 2 kb around peak centers was computed for the merged peak list and shown as a heatmap. Positions in color show high enrichment, and white shows no enrichment. Associated GO terms are shown at right. Oct4 binding data are from ESCs (D0); Oct1 binding data are from ESCs and at days 3 and 6 of mesodermal differentiation. (**D)** Oct1 enrichment based on tag density at peak center at annotated bivalent genes in ESCs (D0), and at MD differentiation days 3 and 6. An additional analysis was performed for MD-specific bivalent genes at MD differentiation D6. (**E)** Example Oct4 and Oct1 binding to three cluster 2 bivalent genes: *Pax3*, *Myog*, and *Pax7*.

We then performed Oct1 ChIP-seq using D3 and D6 MD-differentiated cells. Only 199 Oct4-bound peaks not occupied by Oct1 in pluripotent cells become occupied by Oct1 at D6 of MD differentiation (Fig.4A). In contrast, 807 shared Oct4 and Oct1 peaks in pluripotent cells continued to be bound by Oct1 at D6. Analysis of these peaks using GREAT (genomic regions enrichment of annotations tool) (*26*) identified enrichment for oxidative stress, ribosomal and mitochondrial organization, Notch signaling, and post-implantation development, including somite formation (fig.S4A). Additionally, >6000 peaks become uniquely bound by Oct1 at MD D6 (Fig.4A), and >2300 peaks were uniquely bound at D3 (fig.S4B). We observed significant (∼30%) overlap between Oct1-bound target genes at D6 in parental cells and genes that showed significantly reduced expression in cKO cells at the same timepoint (FDR < 0.05; negative log2(FC)) (Fig.4B).

To analyze the overall pattern of Oct1 and Oct4 occupancy during MD differentiation, we applied hierarchical clustering. We used a union of all peaks (Fig. 4C), as opposed to peak intersect (Fig.4A), to assess varying amounts of Oct1 enrichment over the course of differentiation. Three major clusters of occupied targets were identified, two of which showed transiently decreased or static Oct1 binding over time (Fig.4C, clusters 1 and 2). GO analysis indicated the involvement of these loci in signaling, mitochondrial function, stem cell differentiation, and regulation of Notch and TGF-β signaling (Fig.4C).

Poised developmental genes are frequently associated with simultaneous H3K4me3 and H3K27me3 chromatin marking (*5*). We queried Oct1 binding at these “bivalent” genes during differentiation. To identify such genes, we intersected ESC H3K27me3-and H3K4me3-enriched ChIP-seq peaks from the ENCODE database. This procedure identified 3861 bivalent genes (table S4). We then intersected this dataset with Oct1 ChIP-seq in pluripotent and MD-differentiated parental cells, observing an increase in binding over time (Fig.4D, green and blue lines). A similar analysis at D6 using just MD-specific bivalent genes (generated by intersecting the bivalent gene dataset with MD development GO:0007498, table S5) showed even stronger Oct1 enrichment (Fig.4D, purple line). These findings indicate that Oct1 robustly binds to lineage-specific genes in both pluripotent cells and their differentiated progeny. We noted Oct1 and Oct4 enrichment at three bivalent, silent, and developmentally poised genes encoding MD and myogenic transcription factors (*Pax3*, *Myog* and *Pax7*) (Fig.4E).

Rather than being maintained, Oct1 binding increased with differentiation in cluster 3 (Fig.4C). GO analysis indicated that genes near these peaks are associated with chromatin-modifying activities, including histone acetyltransferase and polycomb group complexes. For example, Oct1 and Oct4 were enriched at cluster 3 genes encoding polycomb complex members (*Ezh2*, *Suz12*, *Ring1* and *Ezh1*) (fig.S5A). In the scRNA-seq UMAP projection, *Ezh2* expression was lower in cKO cells at D6 compared to parental cells at the same time point, in particular in paraxial mesoderm (fig.S5, B and C). In pseudotime, Ezh1 failed to become induced later in development (fig.S5D). These data indicate that during MD differentiation, Oct1 additionally binds to, and mediates the induction of, genes encoding epigenetic regulators associated with H3K27me3.

### Oct1 binds near nodes of chromatin loops in differentiating ESCs

To understand the action of Oct1 at distal regulatory elements, we intersected Oct1 ChIP-seq events in parental, Oct1-sufficient mouse ESCs that are carried forward into MD differentiation with the ENCODE enhancer (H3K27Ac/H3K4me1) dataset (>3100 active ESC enhancers). Many Oct1 binding events were associated with these enhancers (Fig.5A). Inspecting these binding events relative to their target gene transcription start sites, we also observed a gradual increase in proximal promoter binding that was very robust by differentiation day 6 (Fig.5B). Intersecting the Oct1 binding events in ESCs with known ESC super-enhancers (*27*) generated an even more significant overlap, with 47/180 super-enhancers showing Oct1 binding (Fig.5C). GO inspection of this set of Oct1-bound super-enhancers identified multiple terms identified with embryonic morphogenesis, such as “muscle cell differentiation”, as expected (Fig.5D).

**Fig. 5.**
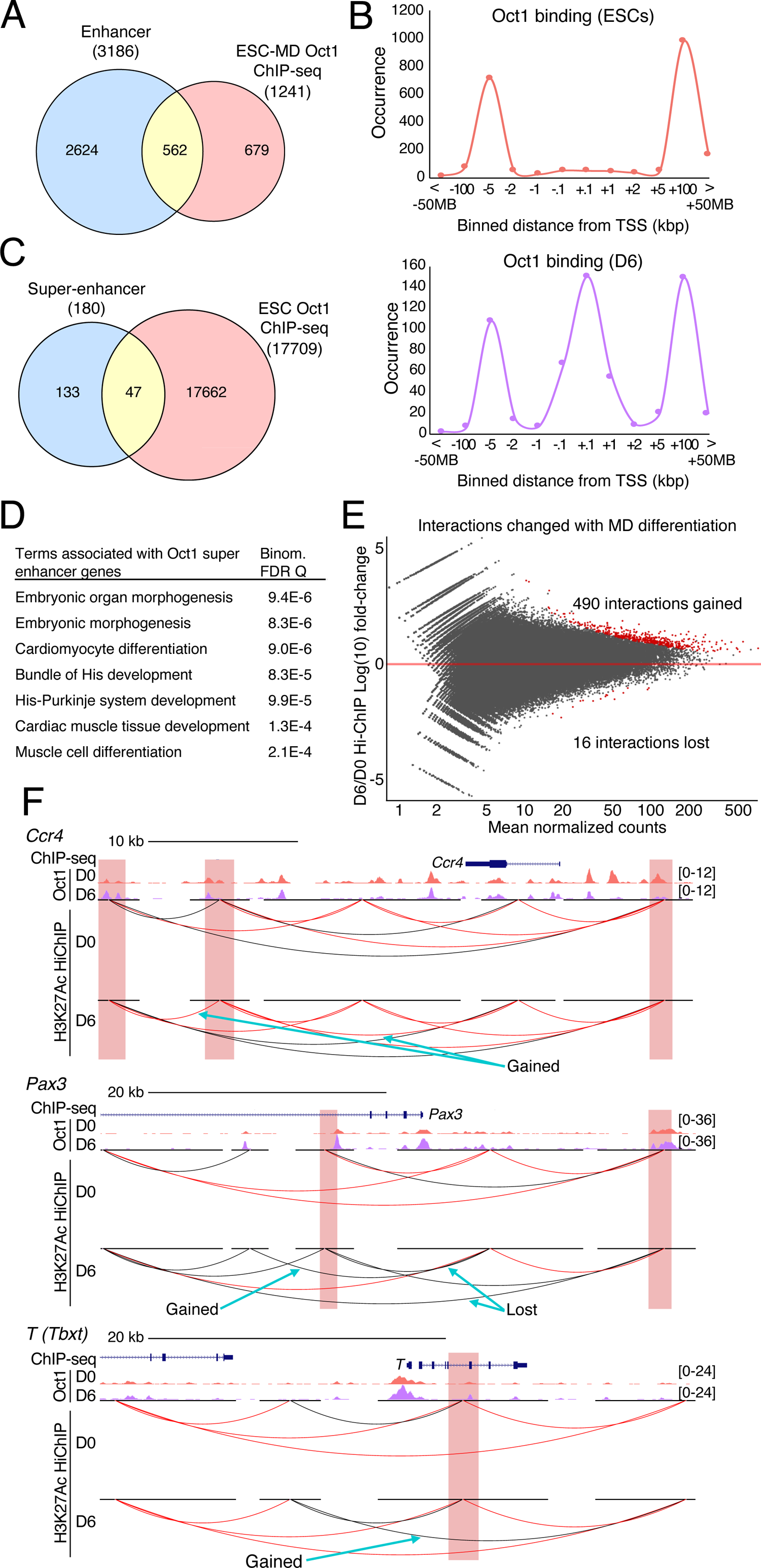
Oct1 occupies developmental enhancer sites at the nodes of topological interactions with target genes. **(A)** Oct1 binding events that were observed in ESCs and that persisted through either D3 or D6 of MD differentiation (ESC-MD Oct1 ChIP-seq) were intersected with known active enhancer regions from ENCODE. Nearly half the peaks (562/1241) were enhancer-associated (0 bp gap). (**B)** Oct1 binding events were mapped to the TSS of the nearest gene and binned by distance. ESCs and MD D6 are shown. (**C)** Annotated super-enhancers (*27*) were intersected with Oct1 binding events in ESCs. Greater than 25% (47/180) were associated with Oct1 binding. (**D)** GO terms associated with Oct1-bound super-enhancers. Binom. FDR Q, Binomial false discovery rate Q-value. (**E)** A scatter plot is shown comparing Log(10) fold-change in H3K27Ac Hi-ChIP interactions and mean normalized counts (2 replicates). D6 differentiated to D0 (pluripotent) parental cells. Adj. *p*-value cutoff ≤0.1 interactions are shown in red. Plot was generated using DEseq2. n = 2. (**F)** Comparison of Oct1 ChIP events with topological interactions in pluripotent parental cells (D0) and after six days of MD differentiation (D6) for three representative genes: *Ccr4*, *Pax3* and *T* (*Tbxt*). For each gene, the group-autoscaled Oct1 ChIP-seq track heights are shown above the *Mm10* H3K27Ac Hi-ChIP genome tracks. Black lines depict 1 to 4 interactions, and red lines 5 or more. Highlighted regions show areas of overlap between TAD anchor sites and Oct1 ChIP-seq binding sites. Highlighted regions were centered ±1 kb over the anchor site.

The increase in promoter binding at D6 suggested that DNA looping may facilitate Oct1 interactions with transcription complexes at proximal promoters. Because no interaction data are available at early steps of MD differentiation, we conducted Hi-ChIP to identify chromatin loops genome-wide. We used pluripotent (D0) and D6-differentiated parental, Oct1-sufficient cells and antibodies targeting the enhancer-enriched H3K27Ac chromatin modification. Two biological replicates were performed for each condition, with between 100M and 200M paired-end reads per replicate. On average, 96.3% of the reads mapped to the reference *Mm10* mouse genome. This procedure produced genome-wide H3K27Ac-enriched topologically-associated domains (TADs) that varied significantly between the two conditions but little between replicates (fig.S6A). We identified 804,599 (D0+D6) chromatin loops, the vast majority of which were maintained after 6 days of MD differentiation (Fig.5E). A small number of interactions (16) were lost; however a 30-fold larger number of interactions were gained (490, Fig.5E).

Comparing the chromatin looping events to the Oct1 ChIP-seq data, we found that the anchor regions of a high proportion of these new interactions corresponded to Oct1 binding events, for both D0 and D6 (fig.S6B). Between 60-70% of the 808 differential interactions (∼500) corresponded to Oct1 binding events using ChIP-seq at D0 and D6. About 10% (93) of the interactions associated with Oct1 binding events were common between D0 and D6 (table S6), such as those for *Ccr4*, *Pax3, T* (*Tbxt*), *Pax7*, *Cd7*, and *Foxk1* (Fig. 5G, fig.S6C). Cumulatively, these results identify a substantial overlap between the termini of chromatin loops and Oct1 binding events early in ESC differentiation to MD.

### Oct1 recruits Utx to *Pax3* to remove H3K27me3

Lineage-appropriate poised developmental genes become activated in part through the removal of repressive H3K27me3 marks (*5, 28*). Our findings indicated that genes with failed induction in differentiating cKO cells tended to be bound by chromatin-modifying complexes that act on H3K27me3 (Fig.1F). We hypothesized that during differentiation, Oct1 locally recruits H3K27-specific demethylases to lineage-specific genes as part of a program that mediates their induction. One such demethylase is Utx, also known as Kdm6a (*29*), which has been shown to promote MD differentiation (*30*). To test the association between Oct1 and Utx binding, we also performed ChIP-seq using H3K27me3 and Utx antibodies at differentiation day 6 using parental, Oct1 sufficient cells. We identified ∼12,000 H3K27me3 peaks and ∼12,000 Utx peaks, each corresponding to ∼11,000 genes with a TSS that lies within 20 kb (table S3). Unsupervised hierarchical clustering together with Oct1 peaks from the same cells identified shared and distinct peaks, including weak Oct1 binding events with strong Utx binding and H3K27me3 association (Fig.6A, cluster 3), and strong Oct1 binding events associated with weaker Utx binding and lack of H3K27me3 (cluster 4). This latter cluster included genes induced in MD such as *Pax3, Pax7*, and *Myog*. We interpret cluster 3 to be developmental genes that remain poised at this timepoint, and cluster 4 to be lineage-appropriate genes that have been derepressed. GO terms associated with these clusters were enriched for developmental and musculoskeletal abnormalities (Fig.6B). Intersecting the Utx and Oct1 ChIP-seq peaks identified a strikingly high degree of overlap, with ∼70% of Oct1-bound peaks also associating with Utx (Fig.6C). A similar fraction of Oct1-bound genes with diminished expression in cKO cells (Fig.4B) were also bound by Utx (916/1293 genes, 71%).

**Fig. 6.**
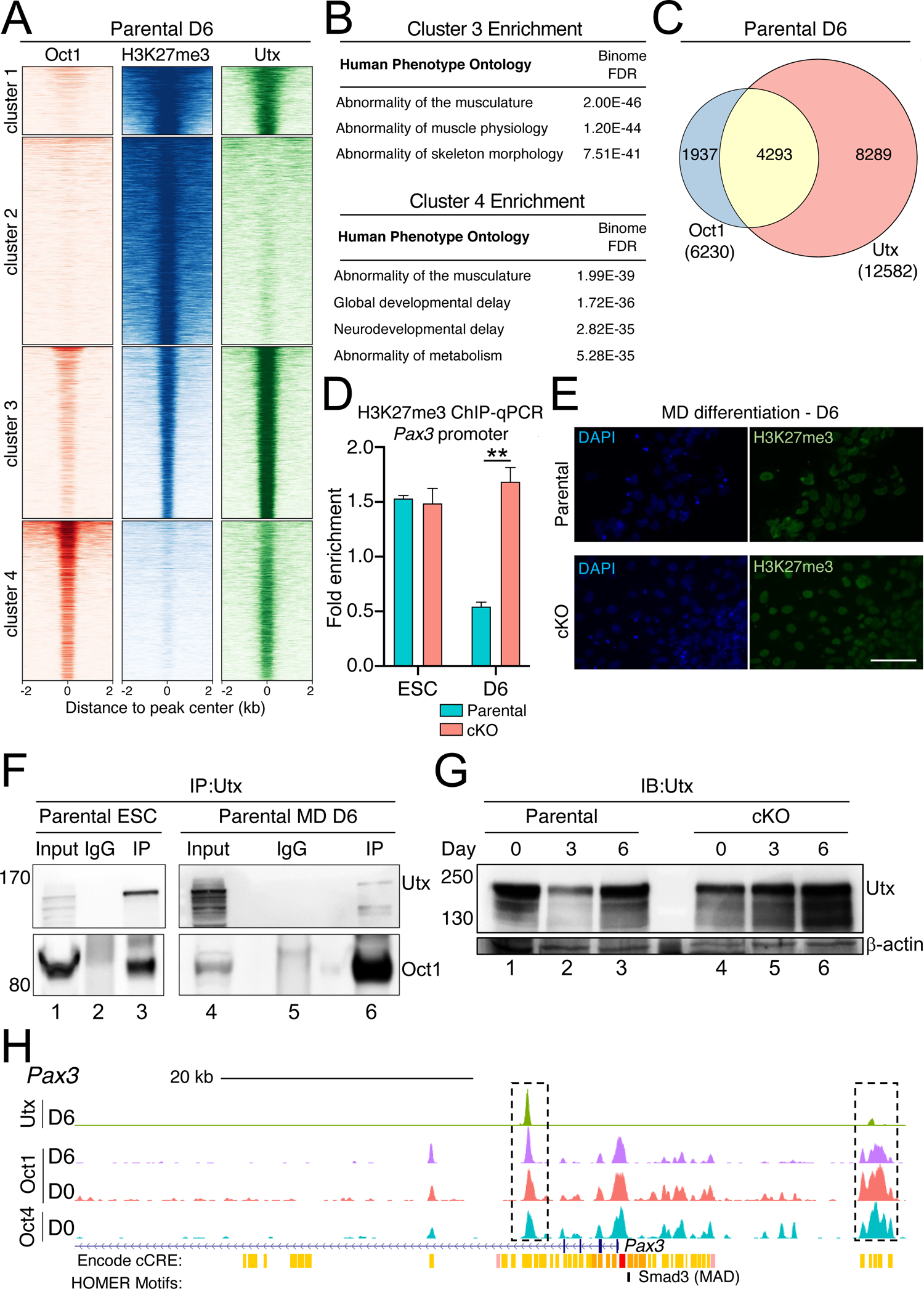
Oct1 recruits Utx to demethylate H3K27me3 at lineage-appropriate genes. **(A)** Oct1, H3K27me3, and Utx ChIP-seq enrichment 2 kb around peak centers was computed for the merged peak list and shown as heatmap based on unsupervised hierarchical clustering. n = 2 biological replicates for all ChIP-seq conditions. (**B)** Human Phenotype Ontology terms for Cluster 3 and 4 genes from (A) are shown. *P*-value was generated using a hypergeometric test. (**C)** The Oct1 and Utx peak lists were intersected (overlap ≥ 1 bp) and plotted as a Venn diagram. 69% of Oct1 peaks overlapped with Utx binding events. (**D)** H3K27me3 ChIP-qPCR enrichment at the *Pax3* promoter in parental and cKO cells at MD differentiation D0 (ESC) and D6. Normalized fold-enrichment is shown relative to both an isotype control antibody and to a nonspecific genomic region encoding the 28S ribosomal subunit. Data represent an average of N=3 biological replicates. Error bars depict ±SEM. (**E)** H3K27me3 immunofluorescence images from D6 MD-differentiated parental and cKO cell cultures. Images were collected a 40× magnification and are representative of 3 independent experiments. Scale bar, 80 μm. (**F)** Utx immunoprecipitation (IP) followed by immunoblotting for Utx and Oct1 using parental ESCs or ESCs MD-differentiated for 6D. Blots are representative of 3 independent experiments. IP with IgG is a negative control. (**G)** Immunoblot (IB) showing Utx in pluripotent (day 0) parental and cKO cells and at days 3 and 6 of MD differentiation. β-actin is a loading control. Blots are representative of 3 independent experiments. (**H)** Signal tracks (*Mm10* v.D191020) showing Oct4, Oct1, and Utx enrichment at the *Pax3* locus 5’ region. Y-axes were group-autoscaled for each gene. Positions of identified HOMER motifs are shown below.

To test if cKO cells inappropriately retained H3K27me3 at a lineage-appropriate gene during MD differentiation, we performed ChIP-qPCR using primers flanking a strong Oct1 peak on the *Pax3* promoter. *Pax3* is induced by D6 of MD differentiation, and its expression is reduced in Oct1 cKO cells at this timepoint (Fig.1E and Fig.3C). As expected, H3K27me3 was robust and equivalent in parental and cKO D0 undifferentiated cells. D6 parental cells showed reduced H3K27me3 at *Pax3*, whereas cKO cells inappropriately retained high H3K27me3 at this locus (Fig.6D). This failure to remove H3K27me3 resulted in ∼3-fold higher H3K27me3 enrichment at *Pax3* in differentiated cKO relative to parental cells (Fig. 6D). Immunofluorescence staining for H3K27me3 showed no difference between parental and cKO cells at D6 (Fig.6E), consistent with there being no substantial global differences in H3K27me3 abundance. These results indicate that both Oct1 and Utx associate with the same sites in *Pax3*—and likely also other genes—and suggest that Oct1 in differentiating cells recruits Utx to locally remove H3K27me3.

We then determined if Oct1 and Utx interact using extracts from undifferentiated and D6-differentiated parental ESCs for coimmunoprecipitation. Oct1 and Utx interacted in ESCs and was maintained during MD differentiation (Fig.6F). Consistently, Utx was present throughout the differentiation timecourse in both parental and cKO cells (Fig.6G). These results support the interpretation that Oct1 interacts with Utx and recruits it to target genes during differentiation to help mediate their activation. Utx recruitment by Oct1 potentially explains the failed induction of lineage-appropriate genes such as *Pax3*, and the failed removal of H3K27me3 in cKO cells (Fig.6D). Oct1 and Oct4 both bound to the *Pax3* promoter in pluripotent cells, whereas Oct1 colocalized with Utx at this locus at D6, early in MD differentiation (Fig.6H).

### Oct1 interacts with Smad3 and cooperates with Smad proteins at *Myog* in cells differentiating towards mesoderm

The broad tissue expression pattern of Oct1 raised the question of how Oct1 mediates gene activation specifically at MD lineage-specific genes. Chromatin accessibility, lineage-specific signaling, and co-bound transcriptional regulators may provide this specificity. We performed motif analysis (*31*) using DNA sequences ±100 bp from the center of the 807 peaks co-bound by Oct1 and Oct4 in pluripotent cells that remain Oct1-bound during MD differentiation (Fig.4A). This procedure identified not only Oct1 motifs, but also motifs for Smads, the terminal transcriptional regulators of intracellular signaling elicited by members of the TGF-β superfamily, including TGF-β, Nodal, and BMP ligands (Fig.7A). TGF-β signals and downstream Smad transcription factor activity are critical for MD induction (*32, 33*). In addition, one study identified Oct1 transcription factor motifs near binding sites for zebrafish Smad2 and Smad3, and further showed that ectopic expression of the zebrafish Oct1 ortholog enhances mesoderm induction, that zebrafish Smad2 and Oct1 physically interact, and that the two factors cooperate to enhance transcription (*34*). Mammalian Smad3 also interacts with Oct4 and co-occupies target genes in pluripotent cells (*35*). Consistent with these findings, Oct1 antibodies efficiently co-precipitated Smad3 in day 6 MD-differentiated cells (Fig.7B). *Smad3* was expressed in undifferentiated parental populations but further induced during MD differentiation (Fig.7C). These findings suggested that Oct1 and Smad3 may co-bind to targets in partially differentiated mesodermal cells. To test this possibility, we intersected the set of Oct1 ChIP-seq targets sites at MD differentiation day 6 with a Smad3 ChIP-seq dataset generated using fibroblasts undergoing early reprogramming into induced pluripotent stem cells (iPSCs) (*36*). We reasoned that this dataset captures cells in a similar state but temporally moving in the opposite direction (dedifferentiation). A high proportion of Oct1 targets (68%) spatially overlapped with these Smad3 target sites (Fig.7D).

**Fig. 7.**
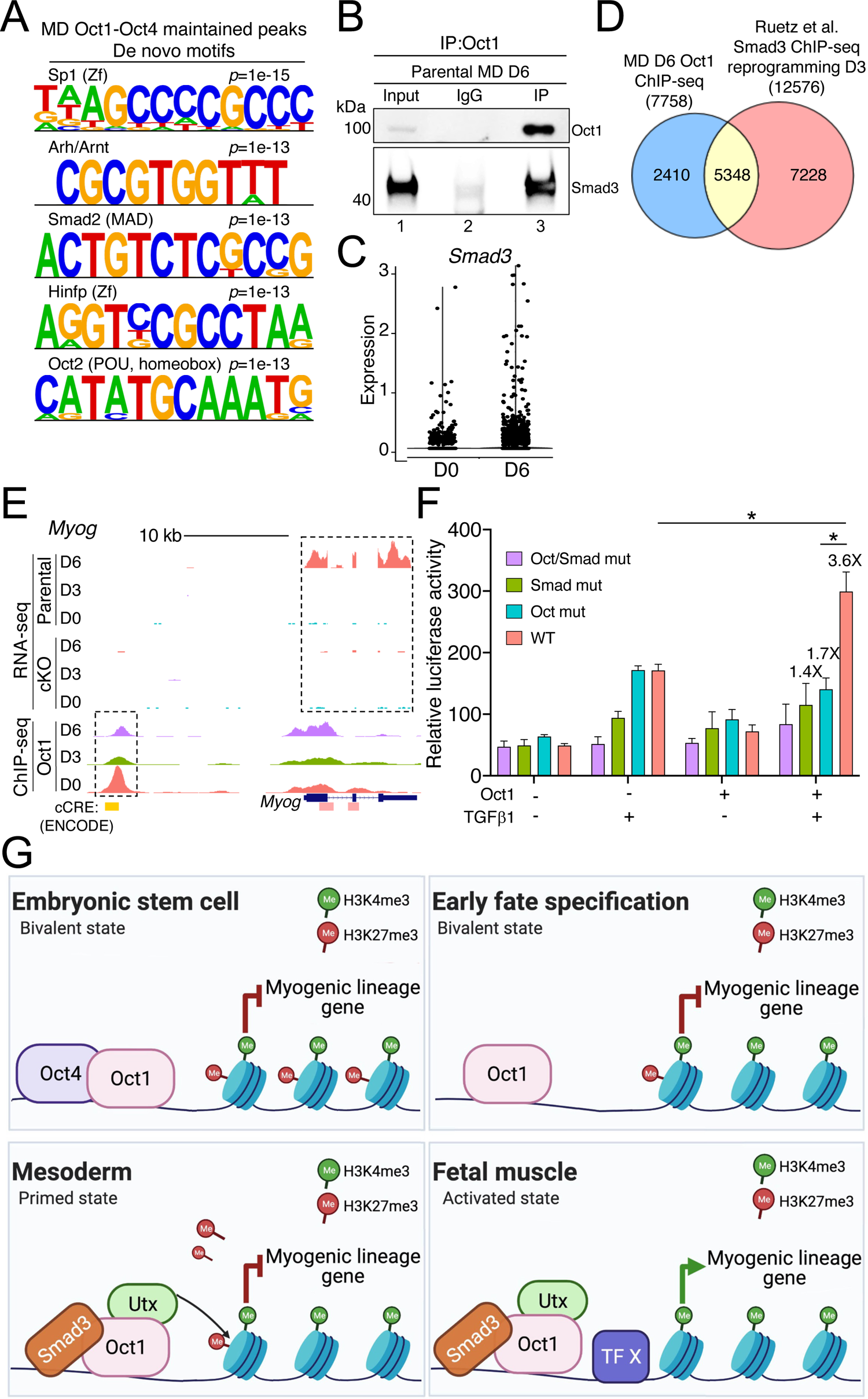
Oct1 and Smad3 cooperate to drive expression of mesoderm-specific genes. **(A)** HOMER de novo motif analysis of Oct1 peaks that are shared with Oct4 in ESCs and maintained after D6 of MD differentiation. The *p*-value output generated by the algorithm indicates the likelihood of false-discovery based on scrambling the nucleotide identities as a control. (**B)** Immunoblotting for Smad3 and Oct1 in D6 MD-differentiated parental cell lysates immunoprecipitated (IP) with Oct1 antibodies, or rabbit IgG as a negative control. Blots are representative of 3 independent experiments. (**C)** *Smad3* expression is shown in violin plots for parental ESCs (D0) and at D6 of MD differentiation, based on the single-cell RNA-seq data. (**D)** Intersection between Oct1 ChIP-seq peaks at D6 of MD differentiation and a published Smad3 ChIP-seq dataset generated using mouse fibroblasts in the process of undergoing reprogramming towards pluripotency (GSE85177_D3_3CA_Dox_Smad3). (**E)** RNA-seq tracks and signal tracks (*Mm10* v. D191020) showing Oct1 ChIP-seq enrichment at the *Myog* locus in undifferentiated (D0) and differentiated (D3 and D6) parental and cKO cells. ENCODE-annotated regulatory elements are shown below. Pink, proximal promoter; yellow, distal enhancer. (**F)** Oct1-deficient MEFs transfected the WT *Myog* enhancer construct or versions of the enhancer construct in which the Oct (Oct mut), Smad (Smad mut), or both (Oct/Smad mut) motifs were mutated were supplied with Oct1, recombinant purified TGF-β1 treatment, or both. Secreted luciferase activity was assessed relative to secreted mCherry expressed from a co-transfected plasmid. An average of N=3 biological replicates is shown. Error bars denote ±SEM. For the situation in which both Oct1 and TGF-β1 were both supplied, fold changes relative to the double-mutant construct are noted. (**G)** Sequential model of bivalency resolution for MD-specific genes. Pluripotent ESCs coexpress Oct1 and Oct4, which both bind to poised targets that are in a bivalent state marked by both activating H3K4me3 and repressive H3K27me3 histone modifications. Early in fate specification, pluripotency and Oct4 are lost, but Oct1 continues to occupy these genes. TGF-β signals prime cells for mesoderm differentiation by activating Smad proteins, which interact with Oct1 specifically at MD-specific targets. Oct1 recruits Utx to these sites, leading to the local removal of H3K27me. Later, other transcription factors serve as primary “on” switches for cell fate–specific gene expression, represented here as fetal muscle. A similar mechanism employing different lineage-specifying transcription factors may restrict Oct1 activity to regulatory sequences appropriate for other lineages.

To gain more insight into mechanism, we tested if Oct1 and Smad proteins synergize to regulate gene expression. The *Myog* gene failed to be induced in D6 differentiating Oct1 cKO ESCs and contained a strong Oct1 binding event overlapping with an annotated distal enhancer ∼3 kb upstream of the *Myog* TSS (Fig.7E). This region is located within a known super-enhancer associated with myotubes (*27*). We cloned an 85-bp region of this enhancer centered on the Oct1 ChIP-seq peak together with the CMV core promoter upstream of a nanoLuciferase reporter cassette. This region contains two octamer sites and two Smad sites, and overlaps with an ENCODE distal enhancer signature (*37*). The reporter vector was transfected into mouse embryonic fibroblasts (MEFs) lacking Oct1 (*17, 38*). The cells were also co-transfected with a plasmid constitutively expressing secreted mCherry (*39*) as an internal standard. To selectively express Oct1 and/or activate Smad proteins, cells were co-transfected with a construct encoding mouse Oct1 and/or treated with recombinant TGF-β1. The two treatments together cooperatively activated the reporter (Fig.7F). For example, constructs containing only active Smad or Oct1 binding sites (Fig.7F, green and blue) displayed little more activity than constructs with no sites (purple), but the two sites together generated significantly increased activity (red). The effect was attenuated in the absence of supplemented Oct1 or Smad activation and recapitulated at one of the strongest Oct1 sites identified in our D6 MD differentiation ChIP-seq dataset, upstream of *Ndufa4l2* and immediately downstream of the skeletal muscle-specific gene *Stac3* (fig.S7, table S3). This region is also located in a site of conservation, overlaps with a known enhancer, and contains multiple binding sites for Oct1 and Smad3. These results indicate that Oct and Smad sites at multiple gene targets can cooperate to drive gene expression.

## Discussion

During development, chromatin transitions from a pluripotent state permissive for all embryonic lineages to a lineage-restricted state. Pluripotent cells maintain genes encoding developmental-specific mediators in a poised configuration that allows for either later induction in lineage-appropriate cell types or later stable repression in all other lineages (*5*). The multifunctional POU domain transcription factor Oct1 is widely expressed. Oct1-deficient animals manifest defective induction of MD-derived somites, cardiac tissue, and blood cells (*16, 17*). The co-expression of Oct1 with Oct4 in pluripotent cells and maintenance of Oct1 expression beyond Oct4 silencing (*15*), together with the severe developmental phenotype of Oct1-null animals, prompted us to study the role of Oct1 during differentiation into the MD lineage. Here, we showed that Oct1-deficient ESCs manifested defective MD differentiation, including failure to generate skeletal muscle myotube structures. These cells failed to express *Myod* and *Myog* mRNAs and myosin heavy chain protein. RNA-seq revealed that Oct1 was essential for induction of the MD lineage-specific gene program, with many lineage-specific genes failing to be properly induced. Undifferentiated Oct1-deficient ESCs increased the expression of *Pou2f3* (*Oct11)* which may reflect Oct1 repression of this gene and/or compensation of Oct1 functions in stress response or metabolism (*25, 38, 40*). Using ChIP-seq, we showed that Oct1 continued to bind the developmental lineage-specific gene enhancers during mesodermal differentiation that it co-bound with its paralog Oct4 in pluripotent cells. Oct1 binding was also gained at the promoters of these genes during differentiation. H3K27me3-specific Hi-ChIP revealed a net increase in topological DNA contacts during mesodermal differentiation and co-localization with Oct1 binding events at many of the new contact sites, indicating that Oct1 tends to utilize new DNA loop interactions as a means of regulating gene expression.

Oct1 helps mediate the activation of many bound lineage-specific genes that are silent and marked by H3K27me3 in pluripotent cells. We showed that Oct1 and Utx binding events significantly overlapped and that Oct1 interacts with Utx, suggesting that Oct1 localizes Utx to MD-specific targets to remove H3K27me3 marks. Specificity is achieved at least in part through cooperative interactions with Smad proteins, known drivers of MD induction. Because Oct1 is widely expressed and results in defective differentiation into multiple lineages (*15*), we anticipate that this mechanism may act widely, with cell type–specificity conferred by the association of Oct1 with lineage-driving transcriptional regulators, such as Smad proteins in the case of MD.

A central role for Oct1 in canalizing differentiating pluripotent cells in early steps of mesodermal specification was shown using scRNA-seq with Oct1 cKO ESCs. In the absence of Oct1, cells undergoing differentiation retained an epithelial state and achieved neuromesodermal and somite-stage gene expression patterns poorly and in reduced numbers. Prior work has associated Oct1 with EMT in a variety of contexts (*41–43*). In scRNA-seq, increased populations of cKO cells appeared and were associated with oxidative stress, consistent with findings that cells lacking Oct1 have increased production of reactive oxygen species (*40*). cKO cells mis-expressed developmentally inappropriate genes (for example, *Epcam* in a neuromesodermal cluster) and underwent inappropriate developmental branching towards poorly differentiated states marked by epithelial gene expression. Unlike the loss of lineage-specific master regulators, wherein cell fate often shifts to a specific alternate lineage, these cells lacked clear lineage signatures. The induction of genes important early in MD differentiation such as *Tbxt* became weaker and lost temporal coherence. Later, there was failed induction of genes such as *Pax7* and *Cxcr4*.

During MD differentiation, Oct1 occupied a subset of genes bound by Oct4 in pluripotent cells. These included lineage-specific developmental mediators and genes encoding chromatin-modifying enzymes. Oct1 occupancy on these latter genes increased with differentiation, suggesting a critical role in canalization of cell fate. Additionally, our data are consistent with a model in which Oct1 recruits the chromatin-modifying enzyme Utx to lineage-specific targets such as *Pax3*. The lack of Utx recruitment to lineage-specific genes in cKO cells is consistent with the abnormal retention of H3K27me3 at the promoter and the failed transcriptional increase observed in the Oct1-deficient condition. No global changes in H3K27me3 were observed, consistent with Oct1-mediated recruitment of Utx to mediate focal removal of these repressive histone marks. An “anti-repression” mechanism for Oct1 involving the removal of repressive chromatin marks has been described for Oct1 previously, for H3K9me2 (*44*). Loss of Utx is compatible with maintainance of an undifferentiated state but interferes with MD differentiation (*28*).

One mechanism that allows Oct1 to gain specificity for MD-appropriate targets is collaboration with Smad3. Smad transcription factor binding sites are enriched near sites of Oct1 binding during MD differentiation. TGF-β signals drive Smad transcription factor DNA binding and are critical for MD specification (*34, 35*). Co-immunoprecipitation experiments using D6 MD-differentiated cells showed an interaction between Oct1 and Smad3, consistent with prior findings in zebrafish (*34*). In our model for Oct1 function at lineage-specific genes during MD specification and later differentiation (Fig.7G), Oct1 and Smad form cooperative complexes at MD-specific genes, and Utx recruitment to Oct1 allows for removal of the repressive H3K27me3 mark, contributing to gene activation. Subsequently, other transcription factors (for example, MyoD) act as primary “on” switches to induce gene expression. Oct1 also binds to and induces the expression of genes encoding PRC complex members, such as *Ezh2*, in parental but not cKO cells. The increased expression of PRC components may solidify lineage specification by aiding the repression of the large number of genes specific to alternative lineages (*45*).

Under MD differentiation conditions, exogenous Oct1 increased expression of the terminal differentiation markers *Myod* and *Myog* while decreasing the early lineage-specification marker *Pax3*, which is transiently expressed during MD differentiation and is necessary for later expression of myogenic genes. Because *Pax3* is not expressed in terminally differentiating cells, these results suggest that ectopic Oct1 may enable transit through a *Pax3*-expressing intermediate to potentiate productive terminal differentiation. More investigation into this pathway is necessary to test if Oct1 can be used to more efficiently differentiate pluripotent cells. Such improvements could have implications for regenerative medicine.

## MATERIALS AND METHODS

### High-throughput datasets

The datasets generated during this study are available through the GEO website [Series record GSE160943]. ENCODE data sets for H3K27me3 and H3K4me3 in ESC-Bruce4 embryonic stem cells are deposited under GSE31039. The published Smad3 dataset used for comparison with Oct1 (*36*) is deposited under GSE85177_D3_3CA_Dox_Smad3.

### Statistical analysis

“Biological replicates” used independently cultured cells, whereas “technical replicates” relied on a later aliquot from a derived component, for example aliquoted RNA for qRT-PCR. No outliers were removed. Differences between groups were analyzed using Microsoft Excel and GraphPad Prism 9 software by two-tailed student T-test. Error bars depict ±standard error of measurement (SEM). For T-test statistical significance, *=*p*-value<0.05, **=<0.01, ***=<0.001.

### Cell culture

Mouse ESCs were cultured as previously described (*46*) with 2i conditions: ERK inhibitor PD0325901 (1 μM, LC Laboratories) and GSK3 inhibitor CHIR99021 (3 μM, LC Laboratories). Cultures were maintained on irradiated feeders (ThermoFisher). Prior to all experiments ESCs were plated on gelatin to deplete the feeders. For Oct1 deletion, cells were treated with 4-hydroxytamoxifen and a completely yellow sub-colony was picked and expanded as published (*15*). For MD differentiation, ESCs were plated on gelatin and cultured as previously described (*20*). Briefly, parental and cKO cells were cultured in N2B27 medium supplemented with recombinant Bmp4 (Peprotech) for 2 d. After 48 hr, media supplementation was changed to RDL (Rspo3, R&D Biosystems; DMSO, MilliporeSigma; LDN193189, Peprotech). Cells were harvested 24 hr (day 3) or 96 hr (day 6) later. For muscle differentiation, media supplementation was switched to HIFL (HGF, Peprotech; IGF-1, Peprotech; FGF-2, Peprotech; LDN193189) and cells cultured for 48 hr (day 8), after which medium was switched to 2% horse serum (ThermoFisher). Cells were harvested on day 11 (for imaging and Oct1 complementation) or day 19 (for RT-qPCR).

### Immunofluorescence

Immunofluorescence was performed as described previously (*47*) with modifications, using rabbit anti-H3K27me3 (MilliporeSigma, ABE44, RRID:AB_10563660) and mouse anti-MyHC-emb (eMyHC, Developmental Hybridoma bank, MF 20-C) antibodies. Secondary antibodies were goat anti-rabbit-Alexa568 (ThermoFisher, A11036, RRID:AB_10563566) and anti-mouse-Alexa568 (ThermoFisher, A11004, RRID:AB_2534072).

### Retroviral Oct1 overexpression

Oct1 was ectopically expressed in ESC cells using a previously described retroviral vector (pBabePuro-Oct1) (*47*). pBabePuro was used as an empty vector control. The vector was co-transfected together with the pCL-Eco packaging plasmid into HEK293T cells to generate retroviral particles. Retroviral supernatants were collected 48 hr later, filtered through 0.45 μm filters and applied to ESCs cultures maintained on puromycin-resistant feeder fibroblasts (ThermoFisher). The mixed population of cells was subjected to selection with puromycin for 48 hr.

### RT-qPCR

RNA was isolated using TRIzol (ThermoFisher). cDNA was synthesized using a SuperScript Vilo cDNA Synthesis Kit (ThermoFisher). RT-qPCR oligonucleotide primers are listed in table S7 and were confirmed to generate a single product of the correct size. To ensure specific PCR amplification, every RT-qPCR run was followed by a dissociation phase analysis (denaturation curve) to confirm the presence of single amplified peaks.

### Bulk RNA-seq

RNA was prepared from three independent cultures of undifferentiated or 3 d and 6 d MD-differentiated parental or cKO ESCs. Poly(A) RNA was purified from total RNA samples (100– 500 ng) with oligo(dT) magnetic beads, and stranded mRNA sequencing libraries were prepared as described using the Illumina TruSeq mRNA library preparation kit and RNA UD Indexes. Molarity of adapter-modified molecules was defined by qPCR using the Kapa Biosystems Library Quant Kit. Individual libraries were normalized to 1.3 nM. Sequencing libraries were chemically denatured and applied to an Illumina NovaSeq flow cell using the NovaSeq XP chemistry workflow. Following transfer of the flowcell to an Illumina NovaSeq instrument, 2×51 cycle paired-end sequencing was performed using a NovaSeq S1 reagent kit. Between 13 and 18 million paired-end reads were generated for each condition. More than 99% of aligned reads mapping to the correct strand.

### Bulk RNA-seq analysis

The Illumina adapter sequence was trimmed using cutadapt version 1.16. Fastq data quality were checked using Fastqc verision 0.11.5. Reads were aligned to the mouse *Mm10* genome using STAR version 2.7.3a in two-pass mode. Aligned reads were checked for quality using the Picard tools’ CollectRnaSeqMetrics command to count the number of read-matching exons, UTRs, introns and intergenic regions, and to calculate normalized gene coverage from the top 1000 expressed transcripts. Between 13 and 18 million paired-end reads were generated for each condition, with >99% of aligned reads mapping to the correct strand. Differentially expressed genes were identified using a 5% FDR with DESeq2 version 1.24.0 (*48*). Genes with a count of at least 50 in one or more samples were tested. Genes showing at least 2.5-fold change of expression at an adjusted *p*<0.01 were selected as differentially expressed. Figures were generated in R version 4.0.0 using functions from ggplots libraries and pheatmap. Gene ontologies were determined using PANTHER (*49*).

### Single-cell RNA-seq

Single cell transcriptomes were analyzed as described previously (*50*). The 10X Genomics Chromium Single Cell Gene Expression Solution with 3’ chemistry, version 3 (PN-1000075) was used to tag individual cells with 16 bp barcodes and specific transcripts with 10 bp Unique Molecular Identifiers (UMIs) according to manufacturer instructions. Briefly, single-cell suspensions were isolated using trypsinization and resuspension in PBS with 0.04% BSA (ThermoFisher). Suspensions were filtered through 40 μm cell strainers. Viability and cell count were assessed using a Countess I (ThermoFisher). For each condition, N=3 biological replicates were combined into a single sequencing run. Equilibrium to targeted cell recovery of 6,000 cells along with Gel Beads and reverse transcription reagents were loaded to Chromium Single Cell A to form Gel-bead-in Emulsions (GEMs). Within individual GEMs, cDNA generated from captured and barcoded mRNA was synthesized by reverse transcription at 53°C for 45 min. Samples were then heated to 85°C for 5 min. Subsequent A tailing, end repair, adaptor ligation and sample indexing were performed in bulk according to manufacturer instructions. The resulting barcoding libraries were qualified on Agilent Technology 2200 TapeStation system and subjected to qPCR using a KAPA Biosystems Library Quantification Kit for Illumina Platforms (KK4842). The multiple libraries were then normalized and sequenced on an Illumina NovaSeq 6000 using the 2×150 PE mode.

### Single-cell RNA-seq data processing and clustering

Sequences from the Chromium platform were de-multiplexed and aligned using CellRanger ver. 3.1.0 (10X Genomics) with default parameters mm10-3.0.0. Clustering, filtering, variable gene selection and dimensionality reduction were performed using Seurat ver.3.1.5 (*51*) according to the following workflow: 1, Cells with <300 detected genes and >10000 genes were excluded further analysis. 2, Cells with <12% UMIs mapping to mitochondrial genes were retained for downstream analysis. 3, The UMI counts per ten thousand were log-normalized for each cell using the natural logarithm. 4, Variable genes (2000 features) were selected using the FindVariableFeatures function. 5, Common anchors between the three parental timepoints (Fig.2A) or parental and cKO D6 (Fig.2C) were identified using FindIntegrationAnchors function that were further used to integrate these sets. 6, Gene expression levels in the integrated set were scaled along each gene and linear dimensional reduction was performed. The number of principal components was decided through the assessment of statistical plots (JackStrawPlot and ElbowPlot). 7, Cells were clustered using a by a shared nearest neighbor (SNN) modularity optimization-based clustering algorithm and visualized using two-dimensional uniform manifold approximation and projection (UMAP). 8, Cluster identities were defined based on the distribution of the specific markers. Differentiational gene expression analysis between the parental and cKO clusters was performed using FindMarkers. Genes with adjusted *p*<0.01 were marked red on scatter plots. Using this analysis, the cell numbers were between 1000 and 2500. Reads/cell ranged between 90,000 and 200,000. Genes/cell ranged between 2,300 and 7,000. The percentage of reads mapping to reference mouse genome was 90% or higher, and the reads confidently mapping to exonic regions was 70% or higher.

### Pseudotime and Velocity analysis

Trajectory analysis of scRNA-seq was performed using Monocle v.2.16.0 (*52*). Parental and cKO sets were filtered using the same parameters as above and merged to generate WT and cKO sets. Cells were ordered based on gene lists for the ESC (beginning) and somite (end) clusters in parental UMAP (Fig.2A). Next, we performed dimensional reduction using the DDRTree method to visualize the dataset, ordered the cells by global gene expression levels, and visualized the trajectory of the dataset. Veolocity analysis was performed using the velocyto package (*53*). Loom files were produced using following paramters: velocyto run10x −m mm10. rmsk.gtf genes.gtf. Gtf files were produced from the Cell Ranger pipeline. Velocity embeddings were produced using the velocyto.r and SeuratWrappers packages. Matrices were filtered using following parameters: nFeature_spliced > 300, nFeature_spliced < 10000, nFeature_unspliced > 200, nFeature_unspliced < 6000, percent.mt < 12. Velocity was calculated using RunVelocity using following paremeters: deltaT = 1, kCells = 25, fit.quantile = 0.02. Velocity embedding were projected on T-SNE maps using the show.velocity.on.embedding.cor function.

### ChIP

ChIP-qPCR and ChIP-seq were performed as previously described (*54*). Briefly, WT and cKO cells were crosslinked with 1% formaldehyde for 10 min and quenched for 5 min using 2.5M glycine. Culture plates were washed using ice cold PBS and cells were harvested by cell scaping. Cells were lysed in Farnham buffer (5 mM Pipes pH 8.0, 85 mM KCl, 0.5% NP-40) and subsequently in RIPA buffer (phosphate-buffered saline, 1% NP-40, 0.5% sodium deoxycholate, 0.1% SDS). Chromatin was sheared using a Covaris sonicator for 5 min (30 sec on/30 sec off) with amplitude=40. Correct chromatin fragmentation was confirmed using 1% agarose gels. 50 μg of chromatin was subjected to IP overnight at 4°C with 4 μg of anti-Oct1 (Novus Biological, NBP2-21584), Oct4 (Santa Cruz, sc-5279, RRID:AB_628051) or H3K27me3 (MilliporeSigma) antibodies. For ChIP-seq, N=2 biological replicates were sequenced separately. For ChIP-qPCR, as a control, we used 5 μg of sheared, non-precipitated input chromatin. Samples were incubated with protein G magnetic beads (ThermoFisher, 10004D) for 5 hr and washed in Low Salt buffer (20 mM Tris-Cl pH 8.0, 150 mM NaCl, 2 mM EDTA, 0.1% SDS, 1% Triton X-100), High Salt buffer (identical but with 500 mM NaCl), LiCl buffer, and Tris-EDTA pH 8.0 plus 1 mM EDTA (TE buffer). Washes were performed at 4°C for 10 min with rotation. For re-ChIP, 2% fragmented chromatin was saved as input and the rest used for IP with the primary antibody (either Utx or Oct1) at 4℃ overnight on a rotator. Samples were then incubated with magnetic beads for 5h at 4℃. The beads were washed for 10 min with Low Salt buffer, High Salt buffer, Low Salt buffer, LiCl buffer, and TE buffer sequentially at 4℃. Chromatin was eluted with 300 µL IP Elution buffer at RT, then diluted 10-fold in RIPA buffer. Diluted samples were then subjected to a second IP with 4 µg of the secondary antibody (Oct1 or Oct4) at 4℃ overnight, and incubated with magnetic beads for 5 hr at 4℃. The beads were washed again as described above, then eluted with 300 µL IP Elution Buffer at RT. Re-ChIP samples, together with the 2% input, were incubated at 65℃ overnight to reverse crosslinking. DNA was purified using phenol-chloroform-isoamyl alcohol extraction followed by PCR clean up. qPCR primers can be found in table S7 and were confirmed to generate a single product of the correct size. The results were reported as qPCR values normalized to input chromatin (gDNA) and non-specific region and presented as fold enrichment.

### ChIP-seq analysis

After chromatin was precipitated as described above, and libraries were sequenced using Illumina NovaSeq. Between 22 and 26 million paired-end Illumina sequencing reads were aligned to the mouse *Mm10* reference genome using Novocraft novoalign v3.8.2, allowing for one random alignment of multi-mapping reads, and providing the adapter sequences for automatic trimming concordant with alignment. ChIP was analyzed using the MultiRepMacsChIPSeq pipeline v12.2, using options “-pe −optdist 10000 −dupfrac 0.15 −mapq 10 −cutoff 2 −tdep 10 −peaksize 500 −peakgap 100 −species mouse −chrskip ‘chrM|PhiX’-blacklist mm10.blacklist.bed”. Intersection of Oct1 with Smad3 target sites used previously published ChIP-seq data (GSE85177_D3_3CA_Dox_Smad3) (*36*).

### Hi-ChIP and analysis

The three-dimensional (3D) genome structure was analyzed using Hi-ChIP, which combines chromosome conformation capture with chromatin immunoprecipitation. Hi-ChIP was performed as previously described (*55*) using H3K27Ac antibodies (Abcam, ab4729, RRID:AB_2118291) and two biological replicates for each condition. The restriction enzyme *Dpn*II was used instead of *Mbo*I. Crosslinked chromatin was sonicated using an EpiShear probe-in sonicator (Active Motif) with three cycles of 30 sec at an amplitude of 40% with 30 seconds rest between cycles. Hi-ChIP libraries were sequenced using a NovaSeq 6000 as paired-end 50 basepair reads to average depth of 300–400 million read-pairs per lane (100-200 million reads per sample). Reads were aligned to the *Mm10* reference genome using HiC-Pro pipeline (*56*) to extract informative unique paired-end tags (PETs). Hichipper (*57*) was used to perform restriction site bias-aware modeling of the output from HiC-Pro and to call interaction loops. These loops were filtered where loops with >1 reads in both biological replicates were kept and used for downstream analyses. Chromatin interactions that significantly change with cellular differentiation were identified using DESeq2 with an adjusted *p*-value cutoff of 0.1. Loop anchors were assigned to the nearest gene using GREAT with a cutoff of 100kb.

### Immunoprecipitation

Cells were lysed with Cell Lysis Buffer (Life Technologies) in the presence of protease inhibitors (EDTA-free tablet, Roche). IP was performed using 500 μg of extract. Extracts were incubated with 4 μg of anti-Utx (Cell Signaling D3Q1I, 33510, RRID:AB_2721244), anti-Oct1 (Novus Biologicals, NBP2-21584) antibodies, or rabbit IgG control (Jackson ImmunoResearch, 011-000-120, RRID:AB_2337123) overnight at 4°C. Protein-antibody complexes were precipitated with protein-G magnetic beads (Thremo Fisher, 10004D) for 3 hr at 4°C with rotation and washed 3 times with Low Salt buffer (20 mM Tris-HCl pH 8.0, 150 mM NaCl, 2 mM EDTA, 0.1% SDS, 1% Triton X-100) plus protease inhibitors. Precipitated proteins were analyzed by immunoblot using anti-Oct1 (Santa Cruz, sc-8024, RRID:AB_628049), anti-Smad3 (Santa Cruz, sc-101154, RRID:AB_1129525) or anti-Utx (Santa Cruz, sc-514859) antibodies.

### Luciferase reporter assay

Single or combination mutations were introduced into the Oct1 and Smad consensus binding sites in the following mouse regulatory elements (*Myog*, *Mm10* chr1:134,285,666-134,285,750, *Stac3*/*Ndufa4l2*, *Mm10* chr10: 127509210-127509477). IDT gBlocks® were synthesized to contain WT sequences or in the case of *Myog* single or combined mutations in the Oct1 or Smad binding sites, fused upstream of the CMV basal promoter (−216-13 relative to TSS). gBlocks were inserted using sequence-and ligase-independent cloning (*58*) upstream of the coding sequence for a secreted nano-luciferase following digestion of vector pNL2.3 (Promega) using *Eco*RV and *Hind*III. The veracity of the cloned inserts was confirmed by Sanger sequencing. Transfections were conducted in 6-well format using Lipofectamine 3000 (ThermoFisher). 200 ng of reporter plasmid were co-transfected into 3T3-immortalized, Oct1-deficient MEFs (*17, 38*) in DMEM media lacking phenol red (ThermoFisher, 31053028, 7666-MB-005) together with 400 ng MMP9-mCherry (*39*) in 1000 ng total transfected DNA. Where indicated, 400 ng pBabePuro-Oct1 was included in the transfection mix. pUC18 plasmid (Jelena’s email indicates “pCL”) comprised the balance of the transfected DNA. Where indicated, transfected cells were provided with 5 ng recombinant mouse TGFβ1 protein (R&D Systems, 7666-MB-005). mCherry fluorescence was determined first by exciting at 570 nm and measuring emission at 610 nm with a 100 msec time delay using a Perkin-Elmer EnVision Xcite Multilabel Plate Reader. Luminescence was measured in 96-well format using Nano-Glo Luciferase (Promega) and a Modulus luminescence plate reader.

## Acknowledgements

We thank G. Kardon and C. Kikani for critical reading of the manuscript. We thank B. Dalley and O. Allen at the HCI High-Throughput Genomics facility and T. Parnell and B. Lohman at the HCI Bioinformatic Analysis Shared Resource for assistance with ChIP-seq and scRNA-seq. We thank Olivier Pourquié from the Harvard Medical School for the assistance with the mesodermal differentiation protocols. We thank D. Picard and D. Wider for the MMP9-mCherry construct. We thank Ac Tan for advice with statistics.

## Author contributions

D.T. conceived the study, and provided administrative and material support. J.P. conceived and supervised experiments, designed experiments and acquired and interpreted data. Y.W., Z.S. and E.P.H. acquired and interpreted data. H.A. analyzed and interpreted data. J.G. provided material support. M.B.C. provided material support, generated reagents and analyzed data. All authors were involved in writing, reviewing and revising the manuscript.

## Data and materials availability

The bulk RNA-seq, single-cell RNA-seq, ChIP-seq, and Hi-ChIP data have been deposited to the NCBI Gene Expression Omnibus (GEO) database (https://www.ncbi.nlm.nih.gov/geo/), series record GSE160943. All other data needed to evaluate the conclusions in the paper are present in the paper or the Supplementary Materials. All cell lines and plasmids are available on request under standard materials transfer agreements with the University of Utah for academic research purposes.

