## supplemental figures 1-7 for "Oct1 cooperates with Smad transcription factors to promote mesodermal lineage specification"

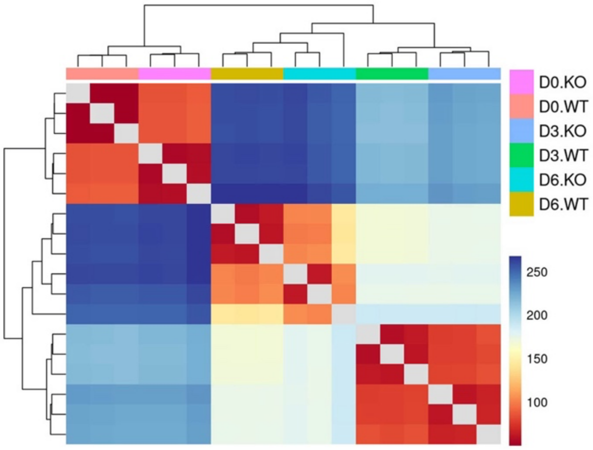


**Fig. S1. Similarity between bulk RNA-seq replicates and conditions.**

Clustering of the RNA-seq replicates using Euclidean distance is shown. D6 shows the most variance compared to the other times and comparing parental (WT) and Oct1 cKO (KO) cells.

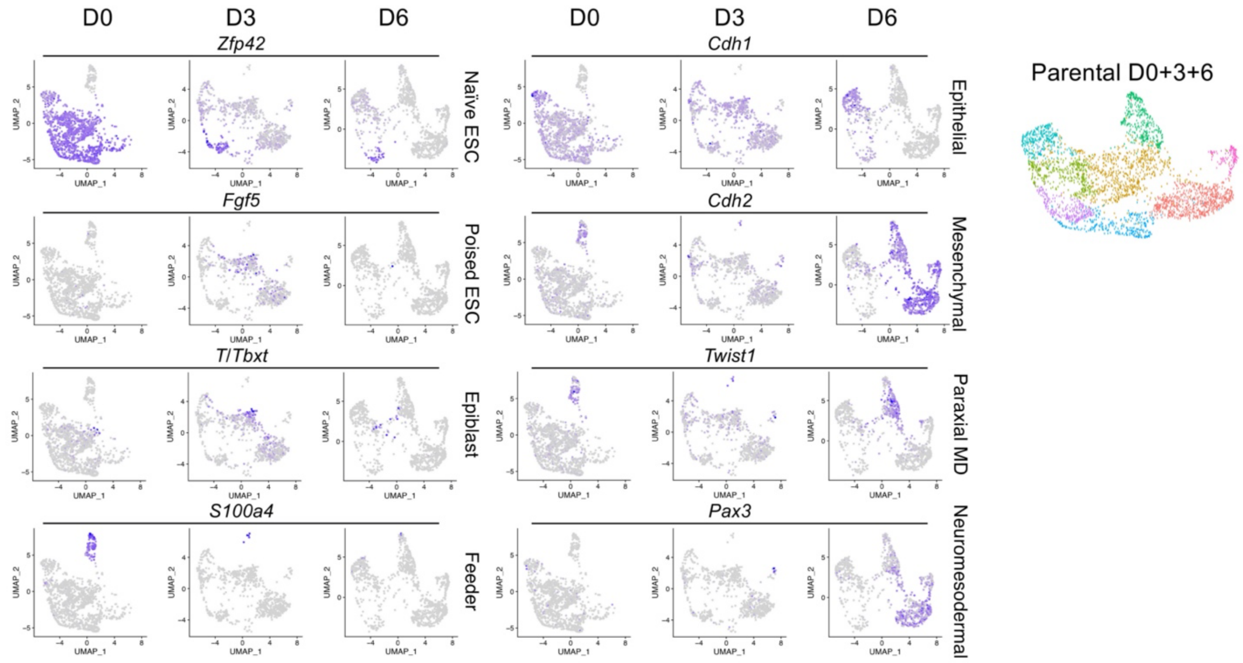


**Fig. S2. Expression of selected genes in UMAP clusters of pluripotent cells and cells at MD differentiation D3 and D6.**

Left: expression of eight different genes specific to different clusters onto a UMAP projection of scRNA-seq data from superimposed parental (Oct1 sufficient) undifferentiated ESCs (D0), and parental cells early during MD differentiation (D3 and D6) used in Fig. 2A. Right: clusters are highlighted in undifferentiated (D0), and in D3 and D6 MD-differentiated cells (similar to Fig. 2A).

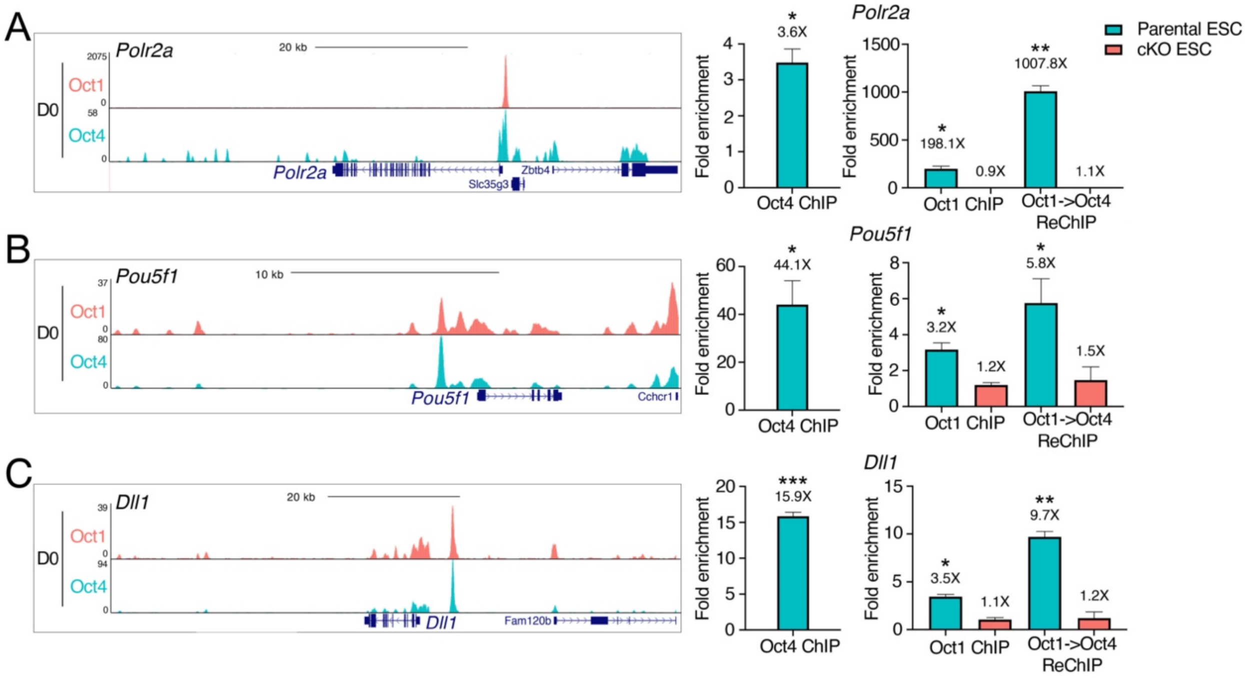


**Fig. S3. Oct1 and Oct4 binding and co-binding in ESC cells (D0 of differentiation).**

(**A** to **C**) ChIP-seq signal tracks for the *Polr2a* promoter (A), *Pou5f1* enhancer (B) and *Dll1* enhancer (C) are shown at left. Plot in center shows Oct1 and Oct4 ChIP-qPCR fold enrichment relative to a nonspecific 40S rRNA genomic region. At far right is Oct1→Oct4 sequential ChIP-qPCR (re-ChIP). qPCR data were normalized to a 40S rRNA nonspecific genomic region. An average of N=3 biological replicates is shown. Error bars depict ±SEM. Student T-test *p*-values: *<0.05, **<0.01, ***<0.001.

A

**D6 Mesoderm**

| GO Biological Process (807 peaks) | Binom FDR *p*-Adj |
|---|---|
| response to oxidative stress | 0.001836177 |
| ribosomal small subunit biogenesis | 0.001804605 |
| mitochondrial membrane organization | 0.014115737 |
| response to reactive oxygen species | 0.020594866 |
| response to hydrogen peroxide | 0.024908306 |
| cellular response to arsenic-containing substance | 0.034804788 |
| regulation of Notch signaling pathway | 0.0362016 |

| GO Cellular Component (807 peaks) | Binom FDR *p*-Adj |
|---|---|
| PRC1 complex | 3.338E-06 |
| cleavage furrow | 0.00029203 |
| cell division site part | 0.0003536 |
| PcG protein complex | 0.00040775 |
| collagen trimer | 0.00768286 |
| cluster of actin-based cell projections | 0.03337456 |
| Golgi stack | 0.04605178 |

| MGI Phenotype (807 peaks) | Binom FDR *p*-Adj |
|---|---|
| embryonic lethality between implantation and somite formation | 8.2454E-05 |
| embryonic lethality between implantation and placentation | 8.9031E-05 |

B

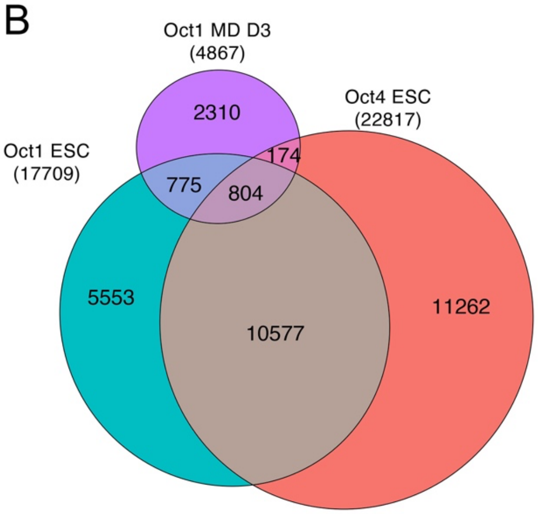


**Fig. S4. Further characterization of Oct1 and Oct4 binding events in 3- and 6-day differentiating parental and cKO ESCs.**

(**A**) Top enriched GO terms in MD D6 Oct1-bound peaks maintained during differentiation (shared with Oct1- and Oct4-bound peaks in ESCs, 807 peaks). (**B**) Venn diagram (similar to Fig. 4A), except showing common and unique binding peaks based on ChIP-seq in parental ESCs (D0) and at D3 rather than D6 of MD differentiation.

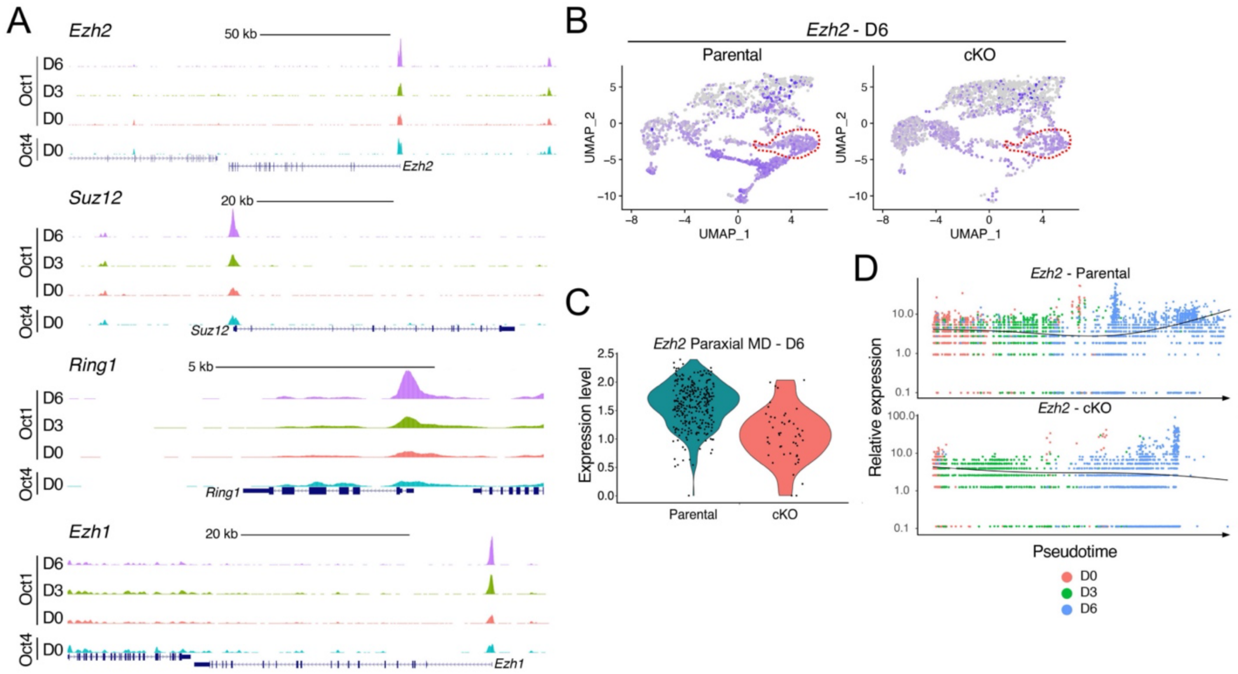


**Fig. S5. Oct1 binding increases at genes associated with regulation of H3K27me3 in ESCs differentiating towards MD.**

(**A**) Representative tracks: *Ezh2*, *Suz12*, *Ring1*, *Ezh1*. Y-axes were scaled to the same value for each gene. (**B**). *Ezh2* expression is shown in UMAP projections for parental and cKO cells at D6 of MD differentiation. The paraxial MD cluster is outlined in red. (**C**). Violin plots showing *Ezh2* expression in cells within the paraxial MD cluster shown in (B). (**D**). *Ezh2* expression in pseudotime in parental (top panel) and cKO (bottom panel) cells. Black trendline represents average expression across all the cells in pseudotime.

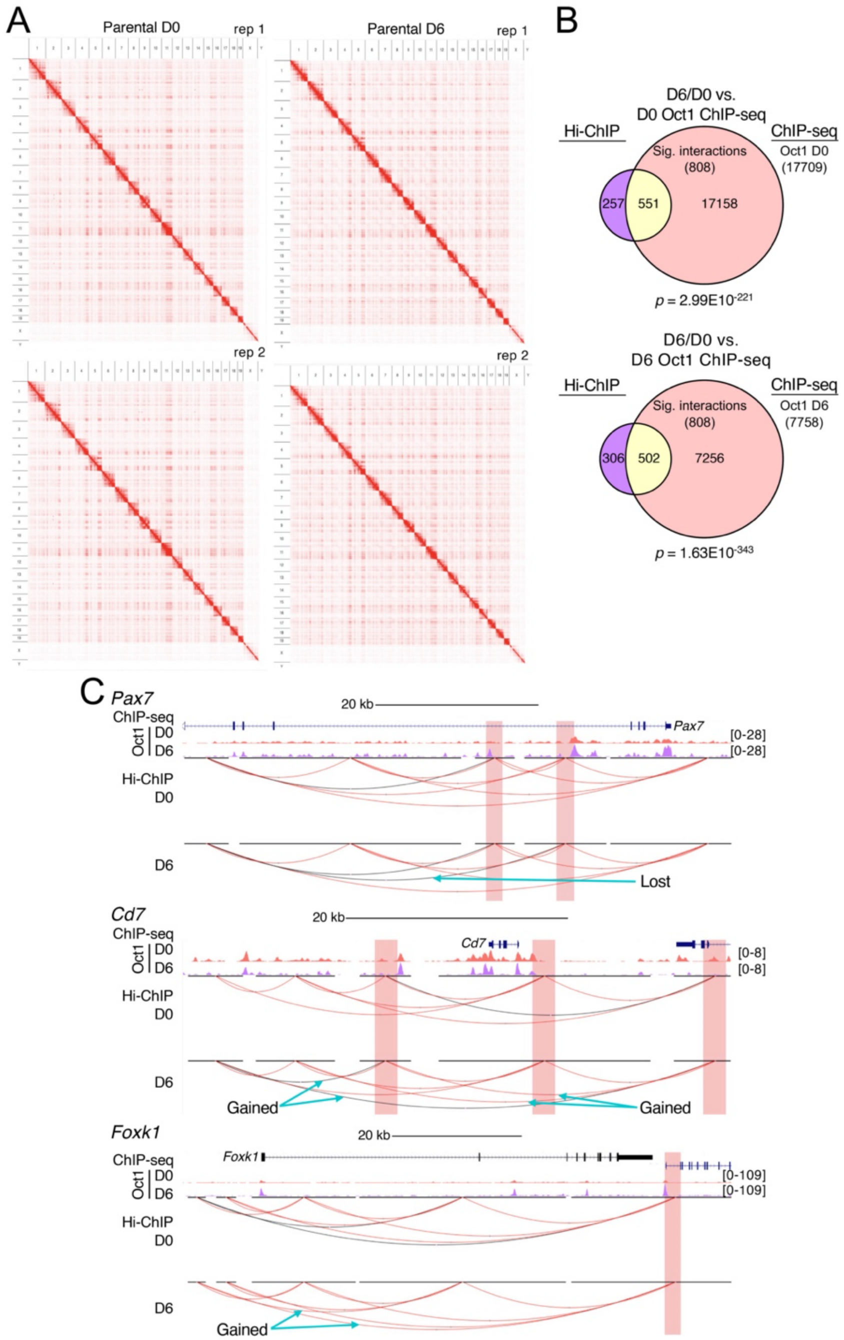
A
Parental D0
rep 1
Parental D6
rep 1
rep 2
rep 2
B
D6/D0 vs.
D0 Oct1 ChIP-seq
Hi-ChIP
ChIP-seq
Sig. interactions
(808)
Oct1 D0
(17709)
257
551
17158
p = 2.99E10^-221
D6/D0 vs.
D6 Oct1 ChIP-seq
Hi-ChIP
ChIP-seq
Sig. interactions
(808)
Oct1 D6
(7758)
306
502
7256
p = 1.63E10^-343
C
Pax7
20 kb
ChIP-seq
Oct1 D0
D6
[0-28]
[0-28]
Hi-ChIP
D0
D6
Lost
Cd7
20 kb
ChIP-seq
Oct1 D0
D6
[0-8]
[0-8]
Hi-ChIP
D0
D6
Gained
Gained
Foxk1
20 kb
ChIP-seq
Oct1 D0
D6
[0-109]
[0-109]
Hi-ChIP
D0
D6
Gained

**Fig. S6. H3K27Ac Hi-ChIP in pluripotent (D0) and six-day differentiated (D6) parental, Oct1-sufficient cells.**

(**A**). Global topologically-associated domains (TADs) represented as a heat map. Replicates were visualized independently. (**B**). Venn diagrams showing overlap between interactions that changed with differentiation and Oct1 binding events. The 506 differential interactions from Fig. 5E were broken down into 808 unique anchor sites showing altered DNA looping, and compared with Oct1 binding at D0. The number of overlapping events represents a 3.85-fold increase over random expectation. The 502 regions at D6 represent an 8-fold increase. *P*-values were computed using a hypergeometric test. (**C**). Similar to Fig. 5F, a comparison of Oct1 ChIP events with topological interactions except using *Pax7*, *Cd7* and *Foxk1* as example genes. Top: group-autoscaled Oct1 ChIP-seq track heights in pluripotent parental cells (D0) and after six days of MD differentiation (D6) for three representative genes: *Ccr4*, *Pax3* and *T* (*Tbxt*). Bottom: *Mm10* H3K27Ac Hi-ChIP genome tracks. Black lines depict 1 to 4 interactions, and red lines 5 or more. Highlighted regions show areas of overlap between TAD anchor sites and Oct1 ChIP-seq binding sites. Highlighted regions were centered ±1 kb over the anchor site.

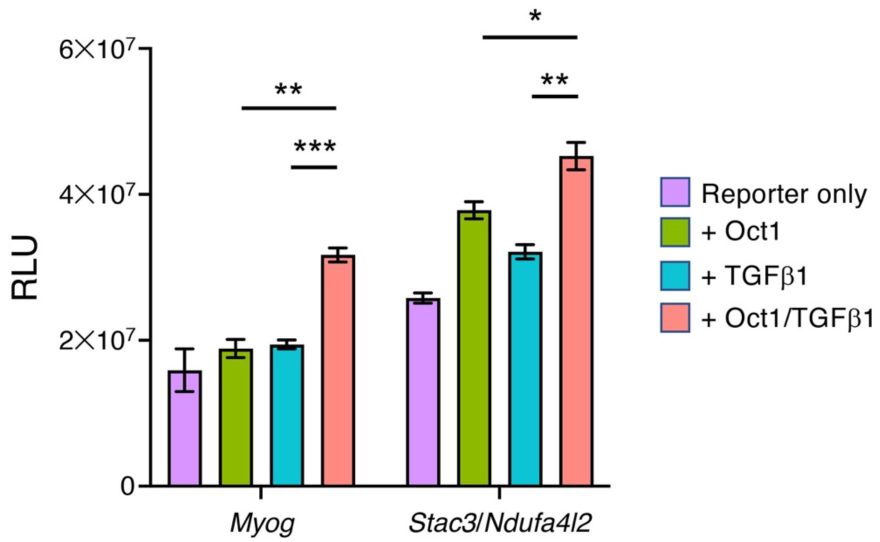


**Fig. S7. Oct1 and Smad2/3 cooperate at an Oct1 target site downstream of the *Stac3* gene to drive luciferase transcription.**

Similar to Fig. 7F, Oct1 deficient MEFs were transfected with reporter constructs and supplied with co-transfected Oct1, recombinant purified TGF-β1, or both. In this case cells were transfected with a reporter construct containing a regulatory DNA sequence corresponding to an Oct1 ChIP-seq binding site identified at MD differentiation D6, located upstream of *Ndufa4l2* and downstream of *Stac3* (Table S3, Oct1 D6 MD, row 5). An average of N=3 biological replicates is shown. Error bars denote ±SEM.

**Table S1. Differential RNAseq for ESC and MD differentiated cells.** Pairwise comparisons in gene expression levels are given for WT (parental) and KO (Oct1 cKO) cells in the pluripotent condition (D0) and at MD differentiation D3 and D6. This table is provided as an Excel file.

**Table S2. Cluster Markers in D6 time point in Parental and cKO integrated set.** For the 14 different clusters (0-13), distinguishing (elevated relative to average levels of the other clusters) are shown. This table is provided as an Excel file.

**Table S3. ChIPseq peaks for Oct4 in ESCs and Oct1 in ESCs, D3, D6 and Utx, H3K4me3 and H3K27me3 at D6 of mesoderm differentiation.** Genomic positions, distance to nearest annotated gene TSS and gene IDs are provided. This table is provided as an Excel file.

**Table S4. Bivalent loci (intersect between GSM769008 and GSM1000089 GEO data sets).** 3861 "bivalent" genes identified by intersecting ESC H3K27me3- and H3K4me3-enriched ChIP-seq peaks from the ENCODE database. This table is provided as an Excel file.

**Table S5. Mesoderm Development GO.** MD-specific bivalent genes identified by intersecting the bivalent gene list with MD development GO:0007498. This table is provided as an Excel file.

**Table S6. Loops with differential interactions on Day 6 versus Day 0 (intersect between GSM769008 and GSM1000089 GEO data sets).** New H3K27Ac Hi-ChIP interactions gained at D6 of MD differentiation were compared to Oct1 binding events. This table is provided as an Excel file.

**Table S7. qPCR primers used in this study.** Oligodeoxynucleotide sequences are provided for both ChIP-qPCR and RT-qPCR. This table is provided as an Excel file.
